# Live-attenuated pediatric parainfluenza vaccine expressing 6P-stabilized SARS-CoV-2 spike protein is protective against SARS-CoV-2 variants in hamsters

**DOI:** 10.1101/2022.12.12.520032

**Authors:** Xueqiao Liu, Hong-Su Park, Yumiko Matsuoka, Celia Santos, Lijuan Yang, Cindy Luongo, Ian N. Moore, Reed F. Johnson, Nicole L. Garza, Peng Zhang, Paolo Lusso, Sonja M. Best, Ursula J. Buchholz, Cyril Le Nouën

## Abstract

The pediatric live-attenuated bovine/human parainfluenza virus type 3 (B/HPIV3)-vectored vaccine expressing the prefusion-stabilized SARS-CoV-2 spike (S) protein (B/HPIV3/S-2P) was previously evaluated *in vitro* and in hamsters. To improve its immunogenicity, we generated B/HPIV3/S-6P, expressing S further stabilized with 6 proline mutations (S-6P). Intranasal immunization of hamsters with B/HPIV3/S-6P reproducibly elicited significantly higher serum anti-S IgA/IgG titers than B/HPIV3/S-2P; hamster sera efficiently neutralized variants of concern (VoCs), including Omicron variants. B/HPIV3/S-2P and B/HPIV3/S-6P immunization protected hamsters against weight loss and lung inflammation following SARS-CoV-2 challenge with the vaccine-matched strain WA1/2020 or VoCs B.1.1.7/Alpha or B.1.351/Beta and induced near-sterilizing immunity. Three weeks post-challenge, B/HPIV3/S-2P- and B/HPIV3/S-6P-immunized hamsters exhibited a robust anamnestic serum antibody response with increased neutralizing potency to VoCs, including Omicron sublineages. B/HPIV3/S-6P primed for stronger anamnestic antibody responses after challenge with WA1/2020 than B/HPIV3/S-2P. B/HPIV3/S-6P will be evaluated as an intranasal vaccine to protect infants against both HPIV3 and SARS-CoV-2.

**AUTHOR SUMMARY:** SARS-CoV-2 infects and causes disease in all age groups. While injectable SARS-CoV-2 vaccines are effective against severe COVID-19, they do not fully prevent SARS-CoV-2 replication and transmission. This study describes the preclinical comparison in hamsters of B/HPIV3/S-2P and B/HPIV3/S-6P, live-attenuated pediatric vector vaccine candidates expressing the “2P” prefusion stabilized version of the SARS-CoV-2 spike protein, or the further-stabilized “6P” version. B/HPIV3/S-6P induced significantly stronger anti-S serum IgA and IgG responses than B/HPIV3/S-2P. A single intranasal immunization with B/HPIV3/S-6P elicited broad systemic antibody responses in hamsters that efficiently neutralized the vaccine-matched isolate as well as variants of concern, including Omicron. B/HPIV3/S-6P immunization induced near-complete airway protection against the vaccine-matched SARS-CoV-2 isolate as well as two variants. Furthermore, following SARS-CoV-2 challenge, immunized hamsters exhibited strong anamnestic serum antibody responses. Based on these data, B/HPIV3/S-6P will be further evaluated in a phase I study.

## INTRODUCTION

SARS-CoV-2 infects and causes disease in all age groups, including infants and children [1]. Even though clinical disease is generally mild in the pediatric population, the overall burden of COVID-19 in this population is substantial [2, 3]. Effective SARS-CoV-2 vaccines are needed for all age groups, including children under 5 years of age. mRNA-based COVID-19 vaccines are available under emergency use authorization for children 6 months to 5 years of age. These and other injectable vaccines are designed to induce strong systemic immune responses [4–6]. However, SARS-CoV-2 is a respiratory virus, which infects and spreads via the respiratory mucosal route. Vaccines with the ability to induce mucosal immunity at the respiratory mucosal entry site are needed [7]. To this end, we are using a live-attenuated bovine/human parainfluenza virus type 3 (B/HPIV3) as a viral vector for intranasal immunization. Recombinant B/HPIV3 was originally developed as a live-attenuated vaccine candidate for intranasal immunization against human parainfluenza virus type 3 [HPIV3] [8, 9]. HPIV3 is an important pediatric respiratory pathogen in humans, second only to human respiratory syncytial virus (RSV) as a cause of severe lower respiratory tract disease in children under 5 years of age. HPIV3 circulates globally, causing annual epidemics, and represents an important vaccine target for the pediatric population [10–12]. The live-attenuated virus B/HPIV3 contains the N, P, M, and L genes of bovine parainfluenza virus (BPIV3), which is attenuated in primates [13]; the genes expressing the parainfluenza virus surface glycoproteins HN and F, the major neutralizing and protective antigens, were derived from HPIV3 [9]. This chimeric live-attenuated vaccine candidate against HPIV3 was immunogenic and well-tolerated in young children [8]. In previous studies, B/HPIV3 was also used as a vector to express the fusion (F) glycoprotein of another human respiratory pathogen, RSV, providing a bivalent HPIV3/RSV vaccine candidate [8, 14]. A version of this vaccine candidate was well-tolerated in children >2 months of age [[15], Clinicaltrials.gov NCT00686075], and improved versions are in further clinical development as bivalent pediatric vaccines against HPIV3 and RSV [16].

In a recent study, we introduced the wildtype and a prefusion-stabilized versions of SARS-CoV-2 spike (S) into the B/HPIV3 vector [17]. The immunogenicity and protective efficacy of these vaccine candidates against a SARS-CoV-2 challenge were evaluated in hamsters, a widely used animal model that supports robust replication of B/HPIV3 vectors and SARS-CoV-2 [18–20]. We found that a single intranasal dose of the B/HPIV3-vectored vaccine expressing the prefusion-stabilized version of SARS-CoV-2 S protein (B/HPIV3/S-2P) induced strong serum antibody responses to SARS-CoV-2 S and elicited serum antibodies with broad neutralizing activity against SARS-CoV-2 of lineages A, B.1.1.7/Alpha and B.1.351/Beta. While a secreted form of the S-2P stabilized version of the SARS-CoV-2 S protein is widely used in injectable mRNA vaccines and vectored or protein vaccines against COVID-19, a further-stabilized secreted version of S was developed, containing 4 additional proline substitutions [S-6P, [21]]. In transient transfection experiments, the expression level of S-6P was about 10-fold higher than that of S-2P, and the overall protein stability and resistance of the S-6P version towards heat stress, storage at room temperature, and multiple freeze-thaw cycles were increased, identifying the secreted version of S-6P as an improved version of S-2P [21].

We recently showed that B/HPIV3/S-6P is immunogenic and protective in rhesus macaques [22]. Here, we extend our work to (i) evaluate the effects of further stabilization by these four additional proline substitutions within the full-length version of the S protein on expression by B/HPIV3, and (ii) to compare the immunogenicity and protective efficacy of B/HPIV3/S-2P and B/HPIV3/S-6P against vaccine-matched SARS-CoV-2 or Alpha and Beta variants in hamsters, and (iii) to compare the breadth of the SARS-CoV-2 antibody response induced by these two S-expressing B/HPIV3 vectors against variants of concern, including three Omicron sublineages.

## RESULTS

### Generation of B/HPIV3/S-6P and comparison of B/HPIV3/S-2P and B/HPIV3/S-6P *in vitro*

We previously used B/HPIV3 to express a prefusion-stabilized version of the full-length SARS-CoV-2 S protein (Wuhan-Hu1, GenBank MN908947), codon-optimized for human expression, and with the S1/S2 polybasic furin cleavage motif “RRAR” ablated by aa substitutions (RRAR-to-GSAS; [23]), resulting in B/HPIV3/S-2P [17]. Here, we generated an additional version, B/HPIV3/S-6P, expressing a full-length version of the SARS-CoV-2 S protein that had been further stabilized by four additional proline substitutions (aa 817, 892, 899 and 942; [21], Fig. 1A). For expression in B/HPIV3, the open reading frames (ORFs) encoding the prefusion-stabilized S proteins were framed by nucleotide adapters containing the BPIV3 gene start and gene end signals for insertion as an additional gene between the BPIV3 N and P ORFs (Fig. 1A).

**Figure 1.**
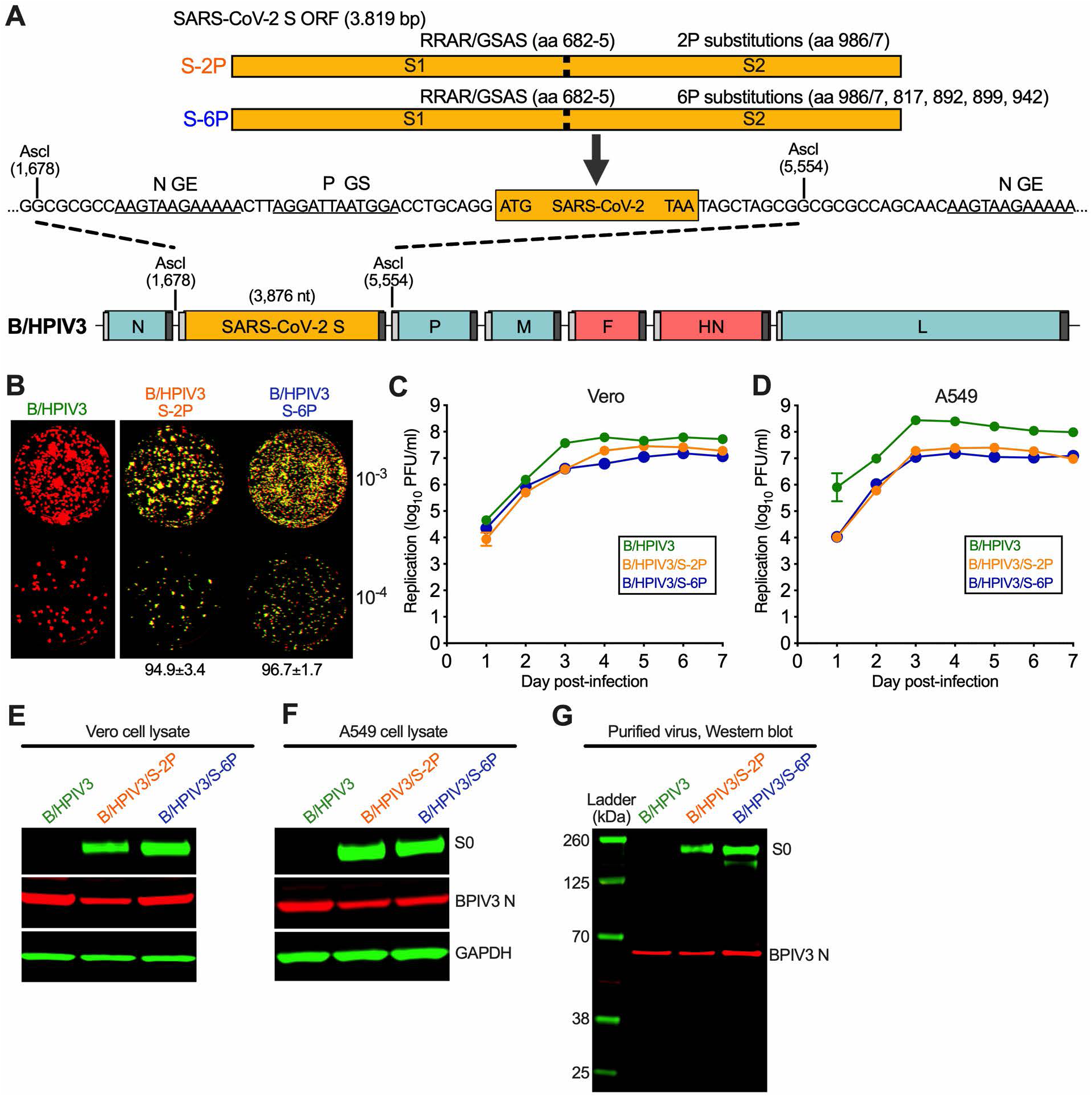
B/HPIV3 vectors expressing prefusion-stabilized versions of the SARS-CoV-2 S protein. (A) Map of the B/HPIV3 genome with the added SARS-CoV-2 S gene. BPIV3 genes are shown in blue, HPIV3 genes encoding the fusion and hemagglutinin-neuraminidase proteins are in red, and the S gene is in orange. Each gene, including the SARS-CoV-2 S gene, begins and ends with PIV3 gene start (GS) and gene end (GE) transcription signals (light and dark grey bars, respectively). The S gene encodes a prefusion-stabilized uncleaved (S-2P) version of the S protein, or a further stabilized version (S-6P) with 6 proline substitutions [6P; [21]], and was inserted into an AscI site between the BPIV3 N and P genes [17]. The stabilizing proline substitutions and four aa substitutions that ablate the furin cleavage site (RRAR to GSAS, aa 682-685) in the S-2P and S-6P proteins are indicated. (B) Stability of SARS-CoV-2 expression, analyzed by dual-staining immunoplaque assay. Virus stocks were titrated on Vero cells and analyzed by dual-staining immunoplaque assay essentially as described [17], using a goat hyperimmune antiserum against a recombinantly-expressed secreted form of S-2P protein and a rabbit hyperimmune antiserum against HPIV3 virions. HPIV3- and SARS-CoV-2 S-specific staining was pseudocolored in red and green, respectively; dual staining appeared as yellow. The percentage (± standard deviation) of yellow plaques is indicated at the bottom. (C, D) Multicycle replication of B/HPIV3 vectors on Vero and human lung epithelial A549 cells. Cells in 6-well plates were infected in triplicate with indicated viruses at a multiplicity of infection (MOI) of 0.01 PFU per cell and incubated at 32°C for a total of 7 days. At 24 h intervals, aliquots of culture medium were collected and flash-frozen for subsequent immunoplaque titration on Vero cells. (E, F, G) Viral proteins in infected cell lysates (E, F) and purified virions (G). Vero (E) or A549 (F) cells in 6-well plates were infected with B/HPIV3, B/HPIV3/S-2P or B/HPIV3/S-6P at an MOI of 1 and incubated at 32°C for 48 h. Cell lysates were prepared and analyzed by SDS-PAGE under denaturing and reducing conditions and Western blotting. The SARS-CoV-2 S protein was detected using the goat hyperimmune serum to S-2P protein, and the BPIV3 N protein was detected by a rabbit hyperimmune serum raised against sucrose-purified HPIV3, followed by immunostaining with infrared fluorophore labeled secondary antibodies and infrared imaging. Immunostaining for glyceraldehyde-3-phosphate dehydrogenase (GAPDH) is shown as a loading control. Images were acquired and analyzed using Image Studio software (LiCor) and are representatives of three independent experiments. (G) Vero-grown virus preparations of B/HPIV3, B/HPIV3/S-2P, and B/HPIV3/S-6P were purified by centrifugation through 30/60% sucrose gradients, and gently pelleted by centrifugation to remove sucrose. One µg of protein per lane was used for SDS-PAGE and Western blotting as described above. Ladder, molecular size marker.

B/HPIV3 expressing the S-6P prefusion stabilized version of the S protein was readily recovered from cDNA by reverse genetics. The sequences of all recombinant B/HPIV3 viruses used in this study were confirmed by Sanger sequencing. We next compared the stability of S expression of S-2P and S-6P by dual-staining immunoplaque assay, performed with polyclonal hyperimmune antisera against PIV3 antigens (green pseudo-coloring), and against the recombinantly-expressed secreted version of the SARS-CoV-2 S-2P protein (red pseudo-coloring). B/HPIV3/S-2P plaques appeared heterogeneous in size, while B/HPIV3/S-6P plaques were more homogenous (Fig. 1B). In virus stocks grown from eight independent recoveries of B/HPIV3/S-2P or B/HPIV3/S-6P, staining for both PIV3 and SARS-CoV-S (yellow, Fig. 1B) was obtained for 94.9±3.4% and 96.7±1.7% of plaques, respectively, indicating that both viruses stably expressed SARS-CoV-2 S protein. In Vero cells, which are widely used for vaccine manufacture, multicycle replication of B/HPIV3/S-2P and B/HPIV3/S-6P was similarly efficient and only slightly delayed compared to B/HPIV3 (Fig.1C), while in human lung epithelial A549 cells, replication of both B/HPIV3/S-2P and B/HPIV3/S-6P was reduced by about 10-fold compared to B/HPIV3 (Fig. 1D).

Next, we compared the expression of the SARS-CoV-2 S by B/HPIV3/S-2P and B/HPIV3/S-6P by Western blot. We infected Vero and human lung epithelial A549 cells at a multiplicity of infection (MOI) of 0.1 plaque-forming unit (PFU) per cell with B/HPIV3 or the S-expressing versions, and prepared cell lysates 24 h after infection. Following SDS-PAGE (under reducing and denaturing conditions) and Western blotting using the hyperimmune antiserum against the secreted version of SARS-CoV-2 S-2P or purified HPIV3 virions, we detected the S protein in lysates at a size consistent with the uncleaved S0 precursor protein (Fig. 1E, F).

We also evaluated the incorporation of the SARS-CoV-2 S-2P or S-6P protein into B/HPIV3 particles. Vero cell-grown viruses were purified by ultracentrifugation through sucrose gradients in three independent experiments. Western blotting revealed that both the S-2P and S-6P versions of the S protein were incorporated in B/HPIV3 particles (Fig. 1G).

### Following intranasal administration, B/HPIV3/S-2P and B/HPIV3/S-6P replicate efficiently in hamsters

To evaluate the replication of B/HPIV3/S-2P and B/HPIV3/S-6P in the hamster model, groups of 28 hamsters were immunized intranasally with 5.0 log_10_ PFU of B/HPIV3/S-2P, B/HPIV3/S-6P, or B/HPIV3 empty vector (Experiment #1, Fig. 2A). On days 3, 5, and 7 post-immunization (pi), 5 hamsters per group were euthanized to evaluate vaccine virus replication in the respiratory tract. Nasal turbinates (NT) and lung tissue homogenates were prepared, and vaccine virus titers were determined by immunoplaque assay (Fig. 2B-C). Additional hamsters from each group were euthanized on days 3 (n=1) and 5 (n=2), and lungs and NT were analyzed by immunohistochemistry (IHC) (Fig. 2D and E). Sera were obtained on day 26 pi from 10 animals per group to evaluate the antibody response.

**Figure 2.**
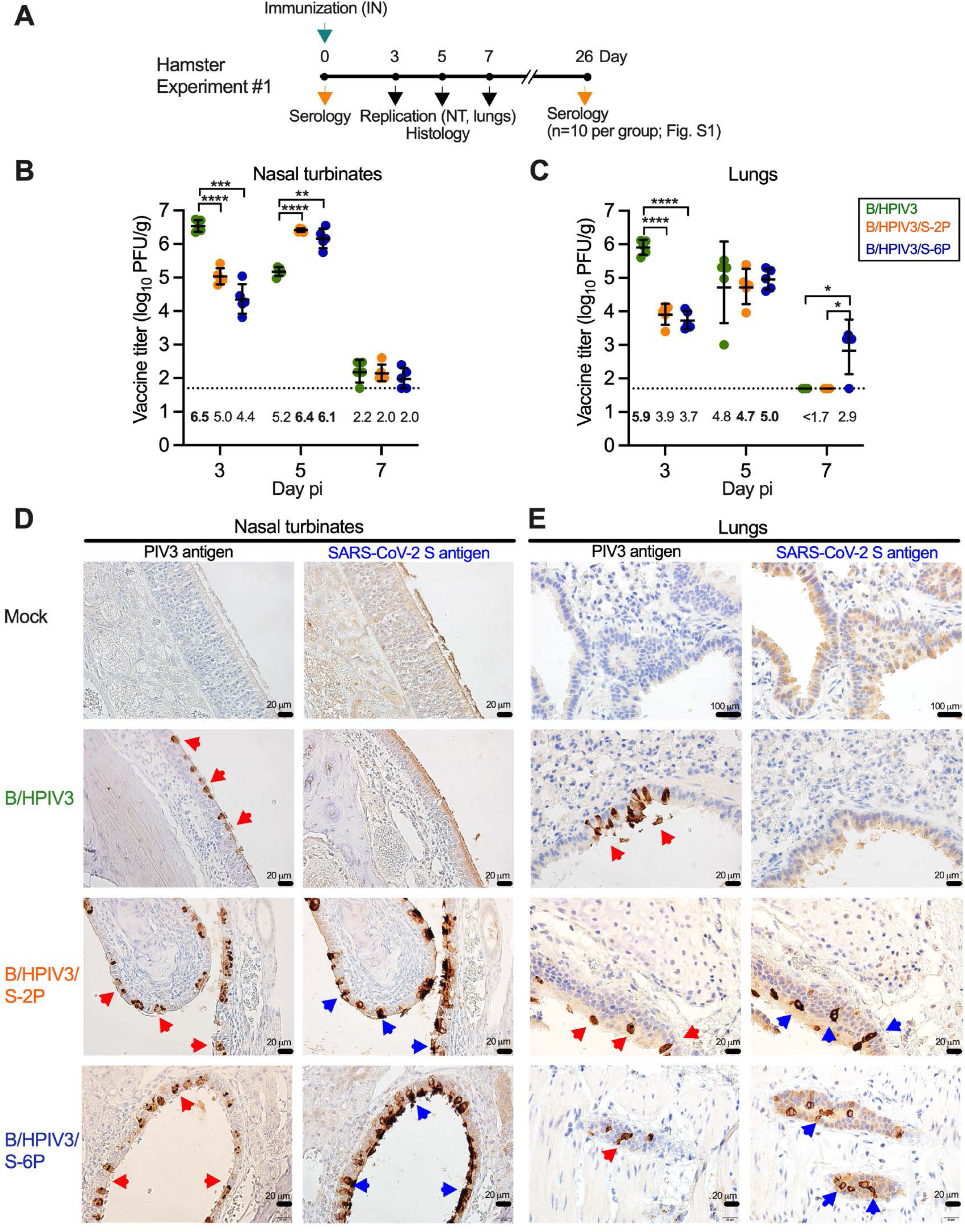
Replication of B/HPIV3, B/HPIV3/S-2P, and B/HPIV3/S-6P in hamsters. (A) In Experiment #1, six-week-old golden Syrian hamsters in groups of 28 were inoculated intranasally with 5 log_10_ PFU of the indicated viruses. On days 3, 5, and 7, five animals per group were sacrificed and the viral titers in the nasal turbinates (NT) (B) and lungs (C) were determined by dual-staining immunoplaque assay. Titers from individual animals are represented by symbols, and geometric mean titers (GMT) and standard deviations are shown by lines. GMT values are indicated below the dotted line; the maximum mean peak titer irrespective of day for each group is in bold. The limit of detection (LOD) was 50 PFU/g of tissue (dotted line). *=P<0.05; **=P<0.01; ***=P<0.001; ****=P<0.0001 (Two-way ANOVA with Tukey multiple comparisons). (D, E) NT (D) and lung tissues (E) obtained on day 5 (n=2 animals per group and n=1 uninfected control animal) were processed for immunohistochemistry. Serial sections were immunostained for HPIV3 and SARS-CoV-2 antigen using hyperimmune antisera raised against HPIV3 virions and a secreted form of the S-2P protein, respectively. Areas with bronchial epithelial cells positive for HPIV3 and SARS-CoV-2 S antigens are marked by red and blue arrowheads, respectively (20 µm or 100 µm size bars are shown in the bottom right corners).

Consistent with our previous results [17], B/HPIV3 replicated well in NT and lungs, with highest titers on day 3 pi (6.5 and 5.9 log_10_ PFU/g, respectively) (Fig. 2B, C). By day 5 pi, B/HPIV3 titers were 20-fold lower in NT, and about 10-fold lower in lungs. B/HPIV3/S-2P and B/HPIV3/S-6P also replicated well. On day 3 pi, titers of B/HPIV3/S-2P and B/HPIV3/S-6P in NT were 30 and 125-fold lower, respectively, compared to the B/HPIV3 empty vector; titers in lungs were 100- and 160-fold lower, respectively, than those of the B/HPIV3 empty vector. By day 5 pi, titers of B/HPIV3/S-2P and B/HPIV3/S-6P had increased in NT (25- and 50-fold, respectively) and lungs (6- and 20-fold, respectively), consistent with delayed replication in the upper and lower respiratory tract due to the presence of the insert. By day 7 pi, replication of B/HPIV3/S-2P, B/HPIV3/S-6P, and empty vector was very low or, in some animals, undetectable in NT or lungs, reflecting self-limiting replication of the vaccine candidates. We also determined the stability of S-2P and S-6P protein expression during *in vivo* replication by dual-staining immunoplaque assay; 96% of B/HPIV3/S-2P and 98% of B/HPIV3/S-6P plaques from NT and 95% of B/HPIV3/S-2P and 99% of B/HPIV3/S-6P plaques from lungs obtained on day 3 after immunization were positive for S protein; 90% and 98% (NT) or 87% and 96% (lungs) of B/HPIV3/S-2P and B/HPIV3/S-6P plaques from specimens obtained on day 5 after immunization were positive for the S protein, showing that expression of both versions of the S protein was stably maintained *in vivo*.

We also evaluated antigen expression in the NT and lungs by IHC. Representative IHC images from NT and lungs obtained on day 5 pi from one of two animals per group are shown in Fig. 2D and E. B/HPIV3 antigen was present in columnar epithelial cells lining the nasal mucosa and the bronchioles in tissue from B/HPIV3, B/HPIV3/S-2P, and B/HPIV3/S-6P-immunized animals (red arrowheads, Fig. 2D, E). SARS-CoV-2 S antigen in animals immunized with B/HPIV3/S-2P and B/HPIV3/S-6P similarly was present in columnar airway epithelial cells of NT and bronchioles (blue arrowheads, Fig. 2D, E). The B/HPIV3 and SARS-CoV-2 S immunostaining patterns and intensities in tissues of B/HPIV3/S-2P and B/HPIV3/S-6P-immunized animals were generally similar. Thus, B/HPIV3/S-2P and B/HPIV3/S-6P efficiently infected and expressed the SARS-CoV-2 S protein in columnar epithelial cells of the nasal and bronchial mucosa.

### B/HPIV3/S-2P and B/HPIV3/S-6P elicited strong serum anti-S and anti-RBD IgG and IgA responses in hamsters

To evaluate immunogenicity and protection against SARS-CoV-2 challenge, 45 hamsters per group were immunized intranasally with 5.0 log_10_ PFU of B/HPIV3/S-2P, B/HPIV3/S-6P, or B/HPIV3 empty vector (Experiment #2, Fig. 3A). We evaluated serum antibody responses 26 or 27 days after intranasal immunization. First, we measured SARS-CoV-2 S-specific serum IgG and IgA (Fig. 3B, C) by ELISA using as antigens the secreted form of the S-2P protein (left panels) and a fragment (aa 319-591) of the S protein bearing the receptor-binding domain (RBD) (right panels). Intranasal immunization with B/HPIV3/S-2P and B/HPIV3/S-6P elicited strong serum anti-S and anti-RBD IgG and IgA responses in hamsters; the geometric mean (GMT) anti-S and anti-RBD IgG titers in the B/HPIV3/S-6P group were significantly higher (1.5-fold, P<0.01, and 2.4-fold, P<0.001) than those induced by B/HPIV3/S-2P. The GMT anti-RBD IgA titers in the B/HPIV3/S-6P group were also significantly higher than those induced by B/HPIV3/S-2P (1.4-fold, P<0.05), while the difference in anti-S IgA titers did not reach the level of significance. We also evaluated the serum antibody response in animals from Experiment #1 (n=10 per group, Fig. S1A), confirming that B/HPIV3/S-6P elicited higher anti-RBD IgG and IgA responses (Fig. S1B, C; P<0.05). In Experiment #1, the differences in the anti-S IgG and IgA responses between B/HPIV3/S-2P and B/HPIV3/S-6P did not reach the level of statistical significance (Fig. S1B, C).

**Figure 3.**
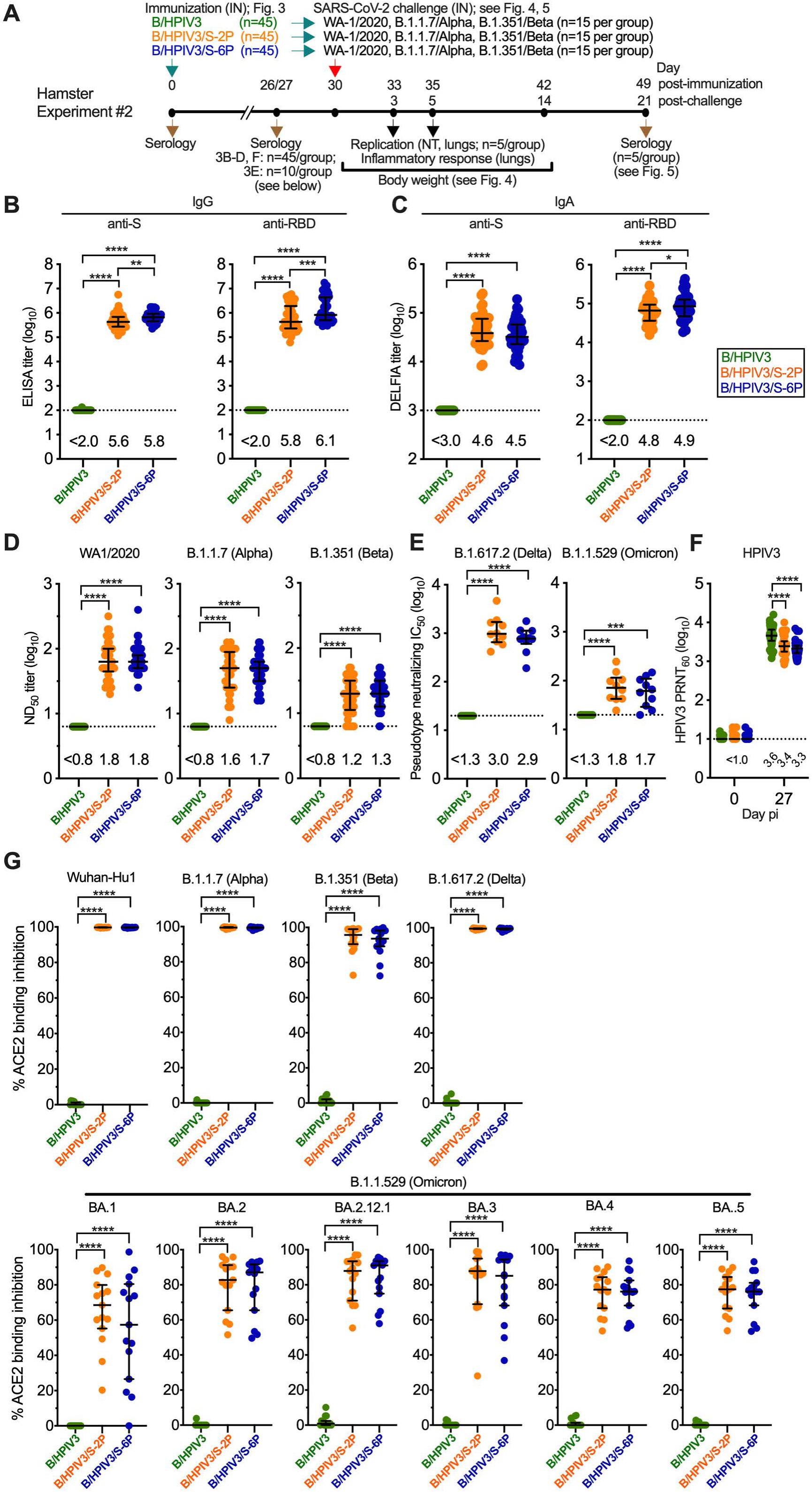
Immunogenicity of B/HPIV3, B/HPIV3/S-2P, and B/HPIV3/S-6P in hamsters. (A) In Experiment #2, six-week-old golden Syrian hamsters in groups of 45 were inoculated intranasally with 5 log_10_ PFU of the indicated viruses. On days 26 or 27, sera were obtained (n=45 per group) to determine IgG ELISA titers to a secreted form of the S-2P protein or to a fragment of the S protein (aa 328-531) containing SARS-CoV-2 receptor-binding domain (RBD) (B). (C) IgA titers to S-2P or the RBD were determined by dissociation-enhanced lanthanide time-resolved fluorescence (DELFIA-TRF) assay. (D) The 50% SARS-CoV-2 neutralizing titers (ND_50_) were determined on Vero E6 cells in live-virus SARS-CoV-2 neutralization assays performed at BSL3 using the vaccine-matched strain WA1/2020, USA/CA_CDC_5574/2020 (B.1.1.7/Alpha variant), and the USA/MD-HP01542/2021 (B.1.351/Beta variant). (E) Ten sera from each group were randomly selected for BSL2 neutralization assays to determine the 50% inhibitory concentration (IC_50_) titers to pseudoviruses bearing spike proteins from SARS-CoV-2 B.1.617.2/Delta or B.1.1.529/Omicron. (F) Sera (n=45 per group) were analyzed to determine the 60% plaque reduction neutralization titers (PRNT_60_) to HPIV3. The detection limits are indicated by dotted lines. (G) ACE2 binding inhibition assay, used as an alternative for a BSL3 live-virus neutralization assay. Heat-inactivated hamster sera were diluted 1:20 and added to duplicate wells of 96-well plates spot-coated with the indicated S proteins. The percent binding inhibition of sulfo-tag labelled ACE2 to S proteins of the indicated SARS-CoV-2 isolates by serum antibodies from immunized hamsters was determined by electrochemiluminescence (for additional antigens, see Fig. S2). Each hamster is represented by a symbol. Medians and interquartile ranges are indicated by lines. *=P<0.05; **=P<0.01; ***=P<0.001; ****=P<0.0001 (One-way ANOVA with Tukey multiple comparisons).

Next, we evaluated the serum SARS-CoV-2-neutralizing antibody responses using the vaccine-matched SARS-CoV-2 strain WA1/2020 (Fig. S1D for Experiment #1 and Fig. 3D, left panel for Experiment #2). We also evaluated virus neutralizing antibody titers to two variants of concern (VoCs; B.1.1.7/Alpha and B.1.351/Beta) in live-virus neutralization assays (Fig. 3D, right panels). B/HPIV3/S-2P and B/HPIV3/S-6P induced robust serum neutralizing antibody titers against all three viruses (Fig. S1D and Fig. 3D). In addition, we randomly selected 10 sera per group to measure antibodies to B.1.617.2/Delta and B.1.1.529/Omicron variants in a pseudotype neutralization assay (Biosafety level (BSL) 2); in both groups, we detected robust B.1.617.2/Delta pseudotype-inhibiting titers; B.1.1.529/Omicron pseudotype-inhibiting titers were about 10-fold lower than those detected in the B.1.617.2/Delta pseudotype assay. As expected, SARS-CoV-2-neutralizing antibodies were not detected in animals immunized with the B/HPIV3 empty vector. We also evaluated the serum neutralizing antibody titers to HPIV3 in animals from Experiment #2 (Fig. 3F) and #1 (Fig. S1E), showing a strong neutralizing immune response in all groups. The B/HPIV3 empty-vector immunized animals exhibited the strongest anti-vector immune response (Experiment #2: 1.6 and 2.1-fold higher than those induced by B/HPIV3/S-2P and B/HPIV3/S-6P, P<0.0001).

Finally, we evaluated the ability of serum antibodies from the B/HPIV3/S-2P- and B/HPIV3/S-6P-immunized hamsters to block binding of soluble angiotensin-converting enzyme 2 (ACE2) to SARS-CoV-2 S proteins of several lineages. This sensitive binding inhibition assay is based on electrochemiluminescence and serves as a high-throughput alternative to BSL3 SARS-CoV-2 neutralization assays. At a serum dilution of 1:20, antibodies from B/HPIV3/S-2P and B/HPIV3/S-6P-immunized animals completely inhibited ACE2 binding to recombinant S proteins derived from the vaccine-matched Wuhan-Hu1 or VoCs B.1.1.7/Alpha and B.1.617.2/Delta (Fig. 3G), and they also efficiently inhibited ACE2 binding to S proteins of B.1.351/Beta or B.1.640.2 (Fig. S2) (93.6 to 97.7 median % inhibition). Furthermore, serum antibodies induced by B/HPIV3/S-2P and B/HPIV3/S-6P also inhibited ACE2 binding to S proteins of the B.1.1.529/Omicron BA.1, BA.2, BA.2.12.1, BA.3, BA.4 and BA.5 sublineages, even though the rate of inhibition was lower (54.1 to 91.1 median % inhibition, Fig. 3G and Fig. S2). No ACE2 binding inhibition was detectable in sera from the B/HPIV3 empty vector-immunized hamsters. Thus, intranasal immunization with B/HPIV3/S-2P and B/HPIV3/S-6P induces a potent neutralizing antibody response with broad activity against SARS-CoV-2 variants, as well as to HPIV3.

### Intranasal immunization with B/HPIV3/S-2P and B/HPIV3/S-6P is protective against challenge with SARS-CoV-2 variants

Hamsters from each of the 3 groups in Experiment #2 (n=45 per group, immunized with B/HPIV3, B/HPIV3/S-2P, or B/HPIV3/S-6P, respectively) were divided randomly into 3 subgroups (n=15 per subgroup). On day 30 after intranasal immunization with B/HPIV3, B/HPIV3/S-2P, or B/HPIV3/S-6P, each of the 3 subgroups were challenged intranasally with 4.5 log_10_ 50% tissue culture infectious doses (TCID_50_) of the vaccine-matched SARS-CoV-2 strain WA1/2020 or VoCs B.1.1.7/Alpha or B.1.351/Beta (n=15 per challenge virus subgroup, Fig. 3A). After challenge with WA1/2020, B.1.1.7/Alpha or B.1.351/Beta, animals previously immunized with B/HPIV3 empty-vector control gradually lost weight over a period of about 7 days (Fig. 4A-C, green lines), and generally started recovering weight beginning on day 7 or 8. Following challenge with WA1/2020, one animal immunized with B/HPIV3 empty-vector control reached the humane study endpoint of 25% weight loss on day 11. On the other hand, hamsters previously immunized with B/HPIV3/S-2P (orange lines) or B/HPIV3/S-6P (blue lines) continued gaining weight after challenge with WA1/2020 and B.1.1.7/Alpha (Fig. 4A, B).

**Figure 4.**
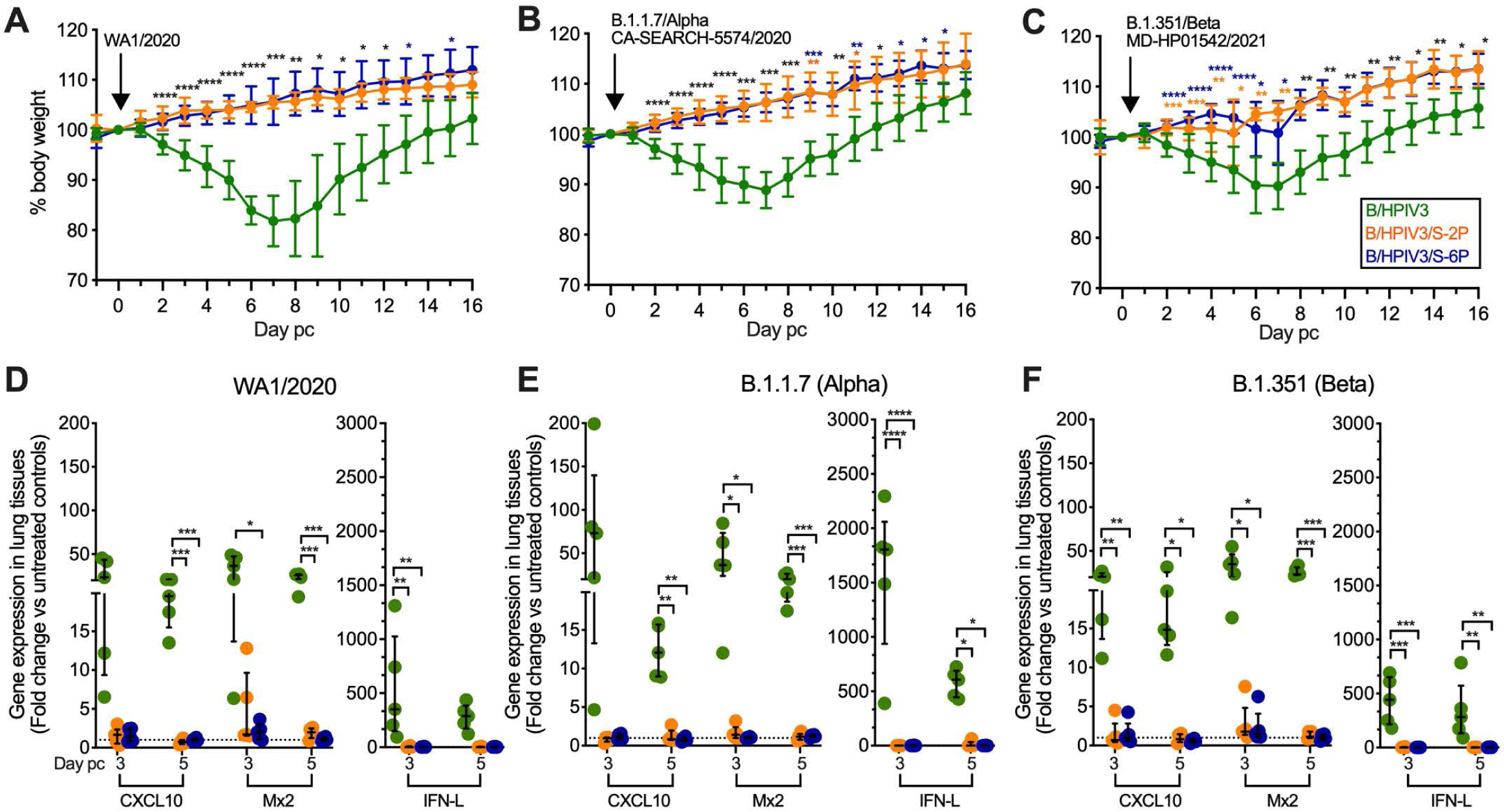
Protection of B/HPIV3, B/HPIV3/S-2P, and B/HPIV3/S-6P-immunized hamsters against weight loss and strong induction of inflammatory cytokines after vaccine-matched and heterologous SARS-CoV-2 challenge. Following immunization of 45 hamsters per group with B/HPIV3, B/HPIV3/S-2P, and B/HPIV3/S-6P in Experiment #2 (see Fig. 3A for schematic), groups were randomly subdivided into challenge groups of 15 animals. On day 30 after immunization, animals were challenged intranasally with 4.5 log_10_ TCID_50_ per animal of the vaccine-matched SARS-CoV-2 strain WA1/2020 (A, D), or representatives of the B.1.1.7/Alpha (B, E), or B.1.351/Beta (C, F) variants. (A-C) Weight changes from day -1 to day 16 post-challenge (pc), expressed as mean % body weight relative to the day 0 time point (n=15 hamsters per group from day -1 to day 3, n=10 hamsters per group on days 4 and 5, and 5 animals from day 6 through 16; in the B/HPIV3 empty-vector control/WA1/2020 challenge group, one animal reached the humane study endpoint of 25% weight loss and was euthanized on day 9; standard deviation of the mean is shown). *=P<0.05; **=P<0.01; ***=P<0.001; ****=P<0.0001 (Mixed-effects analysis with Tukey multiple comparisons). (D-F) Expression of inflammatory cytokines in lung tissues on days 3 and 5 post-challenge. Five animals per group were euthanized on indicated days, and tissues were collected. Total RNA was extracted from lung homogenates and cDNA was synthesized from 350 ng of RNA and analyzed by hamster-specific TaqMan assays. Relative gene expression of C-X-C motif chemokine ligand 10 (CXCL10) and of myxovirus resistance protein 2 (Mx2), a type 1 IFN-inducible antiviral gene, and interferon lambda (IFN-L) compared to the mean level of expression of unimmunized, unchallenged controls (dashed line). qPCR results were analyzed using the comparative threshold cycle (ΔΔC_T_) method, normalized to beta-actin. Each hamster is represented by a symbol. The medians with interquartile ranges are shown. *=P<0.05; **=P<0.01; ***=P<0.001 (Two-way ANOVA with Tukey multiple comparisons).

Following challenge with B.1.351/Beta (Fig. 4C), the weight gain in B/HPIV3/S-2P or B/HPIV3/S-6P-immunized animals was somewhat less uniform between days 2 and 7 after challenge, but the mean body weight in both vaccine groups was significantly higher compared to the corresponding B/HPIV3 empty-vector control-immunized group over this period.

To further evaluate protection from challenge with SARS-CoV-2 WA1/2020, B.1.1.7/Alpha or B.1.351/Beta, we euthanized 5 animals per group on days 3 and 5 after challenge, and extracted RNA from lung homogenates to evaluate the relative expression of inflammatory cytokines C-X-C motif ligand 10 (CXCL10), myxovirus resistance protein 2 (Mx2), or interferon lambda (IFN-L) in response to SARS-CoV-2 challenge by TaqMan assay (Fig. 4D-F)). In previous studies, we had identified CXCL10 and Mx2 as the most strongly induced interferon response genes in hamsters following intranasal inoculation with SARS-CoV-2 [17], serving as biomarkers, together with IFN-L, for the SARS-CoV-2 induced cytokine storm and providing correlates of disease severity [24]. Indeed, after challenge with WA1/2020, B.1.1.7/Alpha or B.1.351/Beta, we detected strong expression of CXCL10, Mx2, or IFN-L in all B/HPIV3 empty-vector control immunized animals (Fig. 4D, E, F). However, after challenge of the B/HPIV3/S-2P or B/HPIV3/S-6P immunized animals with any of these three challenge strains, we found only low or baseline expression of CXCL10, Mx2, or IFN-L, confirming that intranasal immunization with B/HPIV3/S-2P or B/HPIV3/S-6P was protective against cytokine or IFN induction after challenge with the vaccine-matched WA1/2020 strain, or representatives of B.1.1.7/Alpha or B.1.351/Beta variants.

We also determined SARS-CoV-2 challenge virus titers after challenge with strain WA1/2020 or VoCs (B.1.1.7/Alpha or B.1.351/Beta) in tissue homogenates of NT and lung tissues obtained on days 3 and 5 from 5 animals per group (Fig. 5A-C). After challenge of B/HPIV3-empty vector immunized animals with WA1/2020, B.1.1.7/Alpha or B.1.351/Beta variants, we detected high SARS-CoV-2 titers in lungs on both days, with peak GMTs of 8.0 log_10_ TCID_50_ per g (WA1/2020, Fig. 5A, left panel) or 7.4 log_10_TCID_50_/g (B.1.1.7/Alpha and B.1.351/Beta; Fig. 5B, C, left panels). Peak GMTs in NT reached 5.6, 5.8, or 4.9 log_10_TCID_50_/g for WA1/2020, B.1.1.7/Alpha or B.1.351/Beta on day 3 (Fig. 5A-C, right panels). In contrast, B/HPIV3/S-2P and B/HIPV3/S-6P immunized hamsters did not have any WA1/2020, B.1.1.7/Alpha or B.1.351/Beta challenge virus detectable in lungs and NT on days 3 and 5 after challenge, with the exception of 5 animals with detectable virus on day 3 after challenge only; specifically, two B/HPIV3/S-2P-immunized hamsters had WA1/2020 challenge virus detectable in NT, and one hamster immunized with B/HPIV3/S-2P and two hamsters immunized with B/HPIV3/S-6P had B.1.351/Beta challenge virus detectable in both NT and lungs on day 3.

**Figure 5.**
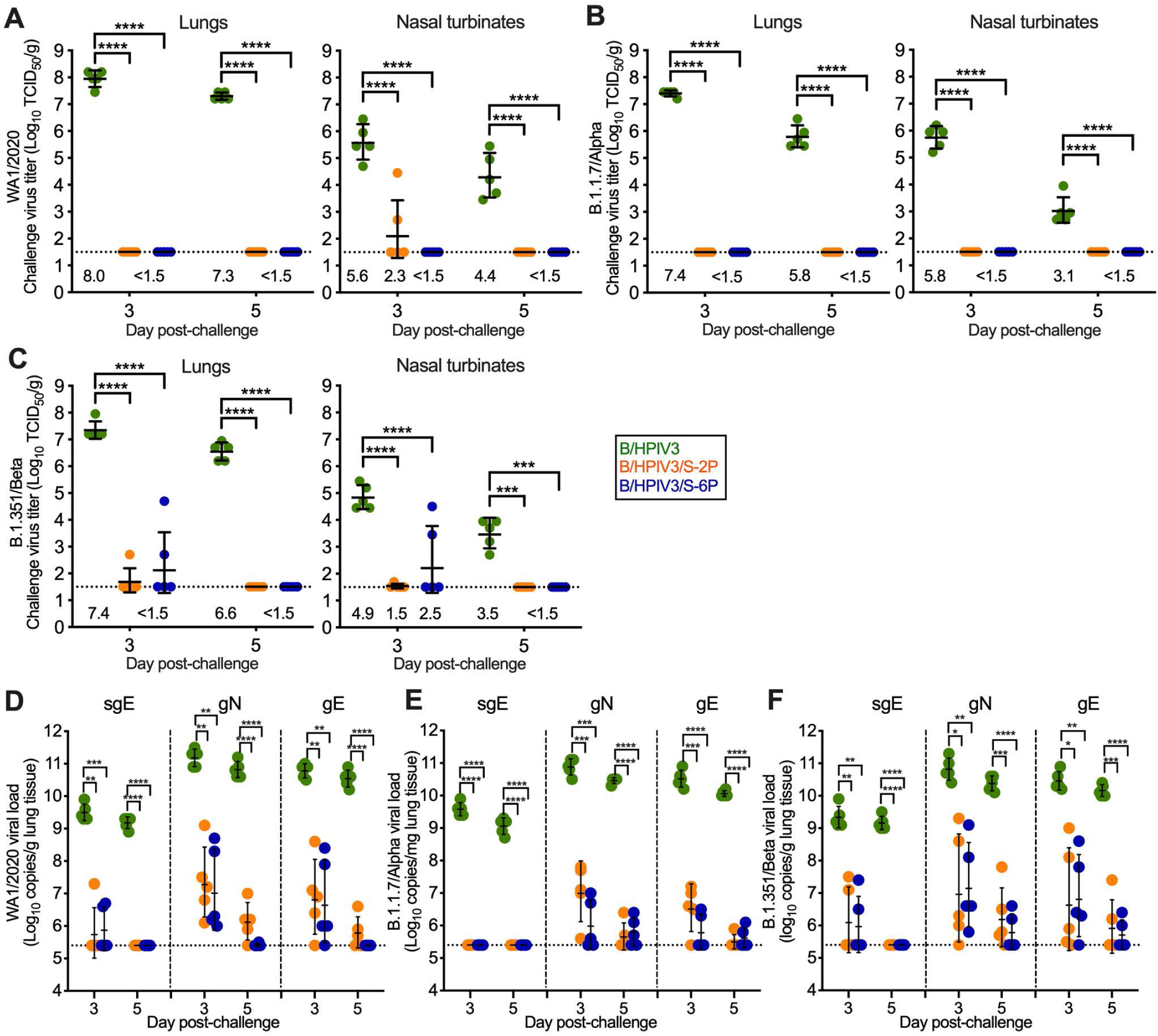
Protection of B/HPIV3, B/HPIV3/S-2P, and B/HPIV3/S-6P-immunized hamsters against SARS-CoV-2 challenge virus replication. As described in Figures 3 and 4, on day 30 after immunization with B/HPIV3 (green symbols), B/HPIV3/S-2P (orange symbols), and B/HPIV3/S-6P (blue symbols), 15 hamsters per group were challenged intranasally with 4.5 log_10_ TCID_50_ per animal of the vaccine-matched SARS-CoV-2 strain WA1/2020 (A, D), or representatives of the B.1.1.7/Alpha (B, E), or B.1.351/Beta (C, F) variants. On days 3 and 5 post-challenge, five animals per challenge virus group were euthanized, and SARS-CoV-2 challenge virus titers were determined in homogenates from lungs (A-C, left panels) and NT (right panels). GMTs are indicated above the x axes. (D-F) SARS-CoV-2 lung viral loads after challenge, expressed in log_10_ genome copies per g. To detect viral subgenomic E mRNA (sgE) RNA, indicative of replicating challenge virus, or genomic N (gN) or E (gE) RNA, indicative of the presence of SARS-CoV-2 challenge virus, cDNA was synthesized using total RNA extracted from lung homogenates as described above, and TaqMan qPCRs were performed (n=5 animals per time point). The GMTs and standard deviations are shown. *=P<0.05; **=P<0.01; ***=P<0.001; ****=P<0.0001 (Two-way ANOVA with Tukey multiple comparisons).

Next, we determined the SARS-CoV-2 viral loads in lungs after challenge. RNA from lung homogenates was evaluated by TaqMan assays to detect genomic N (gN) and E (gE) SARS-CoV-2 RNA, as well as subgenomic E (sgE) RNA. sgE RNA served as an indicator for active replication of challenge virus (Fig. 5D-F). In tissues obtained from all B/HPIV3 empty-vector immunized animals on days 3 or 5 after SARS-CoV-2 challenge, we detected very high viral loads (ranging between 9 and 11 log_10_ copies per g) of gN, gE, and sgE per g of lung tissue. The high levels of sgE copy numbers confirmed high levels of WA1/2020, B.1.1.7/Alpha or B.1.351/Beta challenge virus replication in the empty-vector immunized groups. In the B/HPIV3/S-2P and B/HPIV3/S-6P immunized groups, the levels of genomic RNA was significantly lower on days 3 and 5 after challenge with WA1/2020, B.1.1.7/Alpha or B.1.351/Beta. The levels of subgenomic RNA were also lower or undetectable in the B/HPIV3/S-2P and B/HPIV3/S-6P immunized groups. Specifically, following WA1/2020 challenge, low levels of sgE RNA were only detectable in one B/HPIV3/S-2P and two B/HPIV3/S-6P-immunized animals on day 3 after challenge; after B.1.351/Beta challenge, low levels of subgenomic RNA were detectable in 2 animals per group on day 3. After B.1.1.7/Alpha challenge of B/HPIV3/S-2P and B/HPIV3/S-6P-immunized animals, no sgE RNA was detectable on day 3; by day 5 after challenge, no subgenomic RNA was detectable in any of the SARS-CoV-2 challenge subgroups of B/HPIV3/S-2P or B/HPIV3/S-6P-immunized animals. Thus, intranasal immunization with B/HPIV3/S-2P or B/HPIV3/S-6P was protective in hamsters against challenge virus replication of a vaccine-matched strain of SARS-CoV-2, and two VoCs (Alpha and Beta).

### Magnitude and breadth of anamnestic serum antibody responses in B/HPIV3/S-2P- and B/HPIV3/S-6P-immunized hamsters following SARS-CoV-2 challenge

On day 21 after SARS-CoV-2 challenge, we collected serum from the remaining animals (n=5 per challenge group) and determined the neutralizing serum antibody titers against WA1/2020, B.1.1.7/Alpha or B.1.351/Beta in live-virus neutralization assays performed at BSL3 (Fig. 6A). We found that (i) challenge with WA1/2020, B.1.1.7/Alpha or B.1.351/Beta boosted the WA1/2020 and B.1.1.7/Alpha neutralizing antibody titers of B/HPIV3/S-2P- and B/HPIV3/S-6P-immunized animals (Fig. 3D; Fig. 6A, left and middle panel). (ii) The strongest boost was detected in B/HPIV3/S-6P-immunized, WA1/2020 challenged animals that exhibited three-fold higher anamnestic serum neutralizing antibody titers against WA1/2020 than B/HPIV3/S-2P-immunized animals (P<0.05, Fig. 6A left panel, red label). (iii) WA1/2020 and B.1.1.7/Alpha challenge also boosted the B.1.351/Beta cross-neutralizing titers of the B/HPIV3/S-2P- and the B/HPIV3/S-6P-immunized animals (Fig. 3D; Fig. 6A, right panel); (iv) surprisingly, challenge with the vaccine-matched WA1/2020 virus or a more closely antigenically related virus (B.1.1.7/Alpha) boosted the neutralizing activity to a more antigenically distinct virus, B.1.351/Beta, more effectively than challenge with the antigenically distant B.1.351/Beta virus itself (Fig. 6A, right panel).

**Figure 6.**
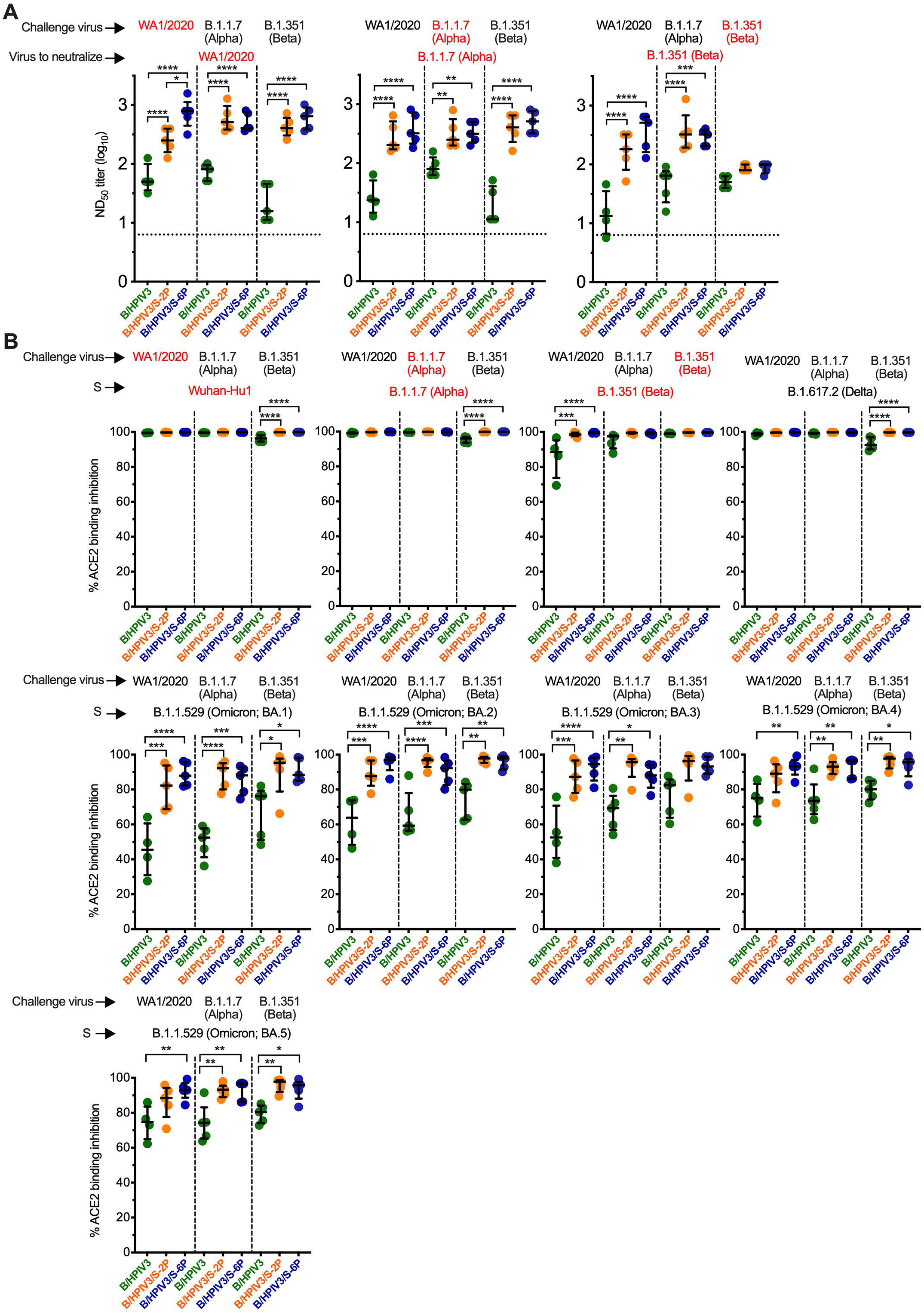
Serum antibody responses in B/HPIV3, B/HPIV3/S-2P, and B/HPIV3/S-6P-immunized hamsters three weeks after challenge with SARS-CoV-2 WA1/2020 or VoCs. As shown in Figure 3A, on day 30 after immunization with B/HPIV3 (green symbols), B/HPIV3/S-2P (orange symbols), and B/HPIV3/S-6P (blue symbols), hamsters were randomly assigned 3 subgroups, and challenged intranasally with 4.5 log_10_ TCID_50_ per animal of the vaccine-matched SARS-CoV-2 strain WA1/2020 or representatives of the B.1.1.7/Alpha or B.1.351/Beta variants. (A) On day 49 (21 days after challenge), sera from 5 animals from each challenge subgroup (shown at the top of the graphs) were collected, and serum neutralizing titers to the SARS-CoV-2 variant indicated in red were determined [WA1/2020 (left graph), B.1.1.7/Alpha (middle graph) or B.1.351/Beta (right graph)]. (B) The same sera were also evaluated in an ACE2 binding inhibition assay, used as an alternative for a BSL3 live-virus neutralization assay. The % ACE2 binding inhibition to ectodomains of S proteins from individual variant (indicated in red) in the presence of serum was calculated relative to ACE2 binding in the absence of serum. Each hamster is represented by a symbol. The median % inhibition and interquartile ranges are shown. The limit of detection is indicated by a dotted line. *=P<0.05; **=P<0.01; ***=P<0.001; ****=P<0.0001 (One-way ANOVA with Sidák’s multiple comparisons).

We also evaluated the ability of the sera from the challenged animals to block binding of a soluble version of ACE2 to S proteins derived from various SARS-CoV-2 lineages, using the ACE2 binding inhibition assay as a high-throughput alternative to BSL3 virus neutralization assays. Twenty-one days after challenge with WA1/2020, B.1.1.7 or B.1.351, sera from B/HPIV3 empty-vector control immunized/SARS-CoV-2 challenged animals inhibited ACE2 binding of S proteins from all 15 lineages and sublineages tested, showing that in the empty-vector control animals, each of the three challenge viruses induced primary antibody responses with broad ACE2 binding inhibition activity to S proteins. Specifically, at the dilution tested, post-challenge sera from B/HPIV3 empty-vector immunized animals completely blocked ACE2 binding to S proteins of the challenged-matched lineages (Fig. 6B, top panels, red labels), and to heterologous WA1/2020 and B.1.1.7/Alpha-derived S proteins (Fig. 6B and Fig. S3, black labels); ACE2 binding inhibition by these sera to the B.1.617.2/Delta-derived S protein also was nearly complete, while the median ACE2 binding inhibition of post-challenge sera from B/HPIV3 control-immunized animals to S proteins of Omicron variants ranged between 45 and 90% (Fig. 6B and Fig. S3).

Sera from B/HPIV3/S-2P and B/HPIV3/S-6P-immunized animals further challenged with WA1/2020, B.1.1.7 or B.1.351 also completely inhibited ACE2 binding to S proteins of WA1/2020, B.1.1.7/Alpha, B.1.351/Beta, B.1.617.2/Delta lineages (Fig. 6B, top row panels), or of the B.1.640.2 lineage (Fig. S3, bottom, right panel). In ACE2 binding inhibition assays to S proteins of all 10 Omicron sublineages tested, WA1/2020, B.1.1.7/Alpha and B.3512/Beta post-challenge sera of B/HPIV3/S-2P and B/HPIV3/S-6P-immunized animals showed significantly stronger binding inhibition compared to post-challenge sera from B/HPIV3 empty-vector control immunized animals, (median ranging between 82 and 99 %, Fig. 6B and Fig. S3), showing that intranasal immunization with B/HPIV3/S-2P and B/HPIV3/S-6P was effective in priming for strong secondary antibody responses with ACE2 binding inhibition activity. Thus, homologous or heterologous SARS-CoV-2 challenge increased the magnitude and breadth of the anamnestic serum antibody responses in B/HPIV3/S-2P and B/HPIV3/S-6P-immunized animals.

## DISCUSSION

Injectable SARS-CoV-2 vaccines provide substantial protection against COVID-19-induced hospitalization and death. However, protection against SARS-CoV-2 infection and shedding, as well as against symptomatic infection, is short-lived, particularly against Omicron variants [25–28]. Ideally, a SARS-CoV-2 vaccine should induce a strong immune response in the upper and lower airways to restrict SARS-CoV-2 replication at the entry site of infection, with the ability to prevent disease. Intranasal vector vaccines will be useful to complement injectable SARS-CoV-2 vaccines for all age groups due to their ability to induce robust protection against SARS-CoV-2 infection and replication in the respiratory tract. The live-attenuated B/HPIV3 vector described in this study has been well-characterized in previous pediatric clinical studies as a pediatric vaccine candidate for intranasal immunization and was safe and immunogenic in infants as young as 2 months of age [8, 15]. In a previous study, we showed that a B/HPIV3 vector expressing the SARS-CoV-2 S protein from the original Wuhan-Hu1 isolate that was stabilized in the prefusion form (B/HPIV3/S-2P) was immunogenic and protective in hamsters [17]. In the present study, we aimed to (i) further improve the immunogenicity of B/HPIV3/S-2P, (ii) evaluate the protection conferred by the S-expressing B/HPIV3 vectors against challenge with SARS-CoV-2 from two heterologous lineages in hamsters, and (iii) evaluate the magnitude and breadth of the antibody response of immunized hamsters following challenge with the homologous or heterologous SARS-CoV-2 isolates.

In our previous study [17], we used the “S-2P” version as SARS-CoV-2 vaccine antigen. This S ORF corresponds to the first available SARS-CoV-2 sequence in which the furin cleavage site “RRAR” was replaced by the “GSAS” motif and two proline substitutions at aa 986 and 987 were introduced to stabilize S into its prefusion form. The resulting B/HPIV3/S-2P was highly immunogenic and protective against a challenge with the homologous SARS-CoV-2 WA1/2020 isolate [17]. However, a subsequent study showed that four additional proline substitutions at aa position 817, 892, 899 and 942 of S introduced into a soluble prefusion-stabilized version of the S protein (S-6P) significantly increased the level of transient expression of S from plasmids in transfected human cells (six to 10-fold), and increased the physical stability of the S protein without affecting its antigenicity [21]. We hypothesized that the potentially increased expression of S-6P compared to S-2P by the B/HPIV3 vector might increase the S-specific immune response. Even though we were not able to detect a statistically significant increase in expression of S-6P over S-2P by B/HPIV3 in Vero or A549 cells, S-6P seemed to be incorporated into Vero-grown B/HPIV3 particles more efficiently than S-2P. Previously, we found that the incorporation of the RSV fusion (F) protein into B/HPIV3 virions increased the magnitude and quality of serum RSV-neutralizing antibodies [16, 29]. We speculated that the increased immunogenicity associated with incorporation into the vector particle could be due to more efficient presentation of the heterologous antigen. In the present study, we found in two independent hamster experiments that B/HPIV3/S-6P induced significantly higher serum anti-RBD IgG and IgA antibody titers than B/HPIV3/S-2P, suggesting that B/HPIV3/S-6P was modestly but significantly more immunogenic than B/HPIV3/S-2P. This might be due to the improvements in the structure-based design, resulting in increased physical stability of the S-6P version [21], and/or the modest increase in incorporation in the B/HPIV3 particles observed in this study.

We also evaluated the breadth of neutralization conferred by the B/HPIV3-S-expressing vectors against VoCs. Serum antibodies from B/HPIV3/S-2P and B/HPIV3/S-6P-immunized hamsters efficiently neutralized the homologous SARS-CoV-2 WA1/2020 isolate as well as VoCs of lineage B.1.1.7/Alpha, B.1.351/Beta or B.1.617.2/Delta. In an ACE2 binding inhibition assay, used as surrogates for BSL3 virus neutralization assays, hamster sera were less efficient in inhibiting ACE2 binding to S derived from B.1.1.529/Omicron sublineages than from WA1/2020 or other VoCs. However, sera from B/HPIV3/S-2P- and B/HPIV3/S-6P-immunized hamsters were still able to substantially inhibit ACE2 binding of S derived from Omicron sublineages. Thus, a single intranasal immunization with the S-expressing B/HPIV3 vectors induced a broad neutralization to antigenically distinct SARS-CoV-2 variants.

B/HPIV3/S-2P and B/HPIV3/S-6P-immunized hamsters were protected from weight loss and lung inflammation following homologous or heterologous (B.1.1.7/Alpha and B.1.351/Beta lineages) SARS-CoV-2 challenge. Only low levels of B.1.351/Beta replication were detected in the NT and lung of B/HPIV3/S-2P and B/HPIV3/S-6P-immunized hamsters that was cleared by day 5 post-challenge. Our results suggested that a single intranasal immunization with B/HPIV3/S-2P or B/HPIV3/S-6P conferred a broad protection in both the upper and lower airways against at least the Alpha and Beta VoCs. Thus, protection induced in the respiratory tract by these vectors seemed to be superior to injectable vaccines such as the adenovirus-based vaccines or alphavirus-based replicating RNA vaccines which in the hamster model were less effective in reducing SARS-CoV-2 challenge virus loads in the lungs [30] or the upper airways of immunized hamsters following challenge with VoCs [31, 32].

Three weeks after challenge, B/HPIV3/S-2P and B/HPIV3/S-6P-immunized hamsters exhibited anamnestic serum antibody responses with an increased breadth of neutralizing activities against all tested SARS-CoV-2 lineages, including 10 different B.1.1.529/Omicron sublineages. Furthermore, B/HPIV3/S-6P appeared to prime for a stronger anamnestic antibody response than B/HPIV3/S-2P, inducing three-fold higher anamnestic neutralizing antibody titers against the homologous SARS-CoV-2 isolate. Since B/HPIV3/S-6P induced more robust primary serum IgA and IgG responses and primed for stronger and broader anamnestic responses, B/HPIV3/S-6P was further evaluated in rhesus macaques [22]. Based on its broad immunogenicity, and the efficacy against two SARS-CoV-2 variants described here, and based on the strong systemic and mucosal immunogenicity in macaques described previously [22], B/HPIV3/S-6P is currently being advanced to a phase I study as an intranasal live attenuated vaccine against both HPIV3 and SARS-CoV-2 for infants and young children.

In this study, we evaluated B/HPIV3 as an intranasal vector vaccine candidate expressing the SARS-CoV-2 S protein from an additional gene. B/HPIV3 is attenuated in humans by the N, P, M, and L genes, which are derived from bovine parainfluenza virus type 3. Thus, B/HPIV3 is expected to be safe in HPIV3 seronegative infants and children and suitable as a bivalent vaccine against HPIV3 and SARS-CoV-2, but it will likely be over-attenuated in HPIV3 experienced individuals. In the best-case scenario, intranasal immunization of HPIV3-experienced individuals with B/HPIV3 expressing the prefusion-stabilized SARS-CoV-2 S antigen would confer a protective mucosal immune response to SARS-CoV-2 due to the high level of immunogenicity of the prefusion-stabilized S protein, and despite the greatly restricted ability of HPIV3 to replicate in the respiratory tract in presence of pre-existing anti-vector immunity. We are planning to evaluate this in prime-boost studies in animal models. In another limitation, in this study, we characterized the differences in prefusion stabilization of two S antigens derived from the ancestral Wuhan Hu-1 isolate, but the currently circulating SARS-CoV-2 variants have evolved to accumulate antigenic differences from the ancestral strain. However, our results show that the breadth of protection and antibody reactivity to antigenically distant variants are substantial. We are currently including S antigens from other variants in our studies to characterize the differences in immune responses and protective efficacy of vaccine-matched vs non-matched antigens in the hamster challenge model.

## MATERIALS AND METHODS

### Cell lines

Human lung epithelial A549 cells (ATCC CCL-185) were grown in F-12K medium (ATCC) with 5% fetal bovine serum (FBS). LLC-MK2 rhesus monkey kidney cells (ATCC CCL-7), African green monkey kidney Vero cells (ATCC CCL-81), or Vero E6 cells (ATCC CRL-1586) were grown in OptiMEM (Thermo Fisher) with 5% FBS. Vero E6 cells were used for SARS-CoV-2 neutralization assays, and to titrate the SARS-CoV-2 challenge viruses.

### Viruses

The SARS-CoV-2 USA-WA1/2020 challenge virus stock was grown on Vero E6 cells stably expressing human TMPRSS2 [17]. The USA/CA_CDC_5574/2020 isolate (lineage B.1.1.7/Alpha variant, GISAID: EPI_ISL_751801; sequence deposited by CDC, isolate obtained from CDC) and the USA/MD-HP01542/2021 isolate (lineage B.1.351/Beta variant, GISAID: EPI_ISL_890360; sequence deposited by Morris et al., Johns Hopkins University (JHU), isolate obtained from Andrew Pekosz, JHU, Baltimore, MD) were also passaged on Vero E6 cells stably expressing TMPRSS2 [17]. Titration of SARS-CoV-2 was performed by determination of the TCID_50_ in Vero E6 cells [33]. All experiments with SARS-CoV-2 were conducted in Biosafety Level (BSL)-3 containment laboratories approved for use by the US Department of Agriculture and CDC.

Virus stocks of recombinant B/HPIV3 [8, 9] and B/HPIV3 vectors were propagated on Vero cells at 32°C and titrated by dual-staining immunoplaque assay [17].

### Dual-staining immunoplaque assay

Briefly, Vero cells in 24-well plates were inoculated with 10-fold serially diluted samples. Two hours later, monolayers were overlaid with culture medium containing 0.8% methylcellulose and incubated at 32°C. Six days later, monolayers were fixed with 80% methanol, and immunostained with an HPIV3-specific rabbit hyperimmune serum to detect B/HPIV3 antigens, and a goat hyperimmune serum against the secreted version of SARS-CoV-2 S-2P to detect co-expression of the S protein. An infrared-dye conjugated donkey anti-rabbit IRDye680 IgG and a donkey anti-goat IRDye800 IgG were used as secondary antibodies. Stained plates were scanned with the Odyssey infrared imaging system (LiCor). Fluorescent staining for PIV3 proteins and SARS-CoV-2 S was visualized in green and red, respectively, generating yellow plaque staining when merged [17].

### Multicycle replication of B/HPIV3 vectors in cell culture

Sub-confluent Vero and A459 cells in 6-well plates were infected in triplicate with indicated viruses at a multiplicity of infection (MOI) of 0.01 PFU per cell. After adsorption for 2h, cells were washed to remove the virus inoculum, and 3 ml of fresh medium was added. Cells were incubated at 32°C for 7 days. Every 24 h, 0.5 ml aliquots of culture medium were collected and snap-frozen in dry ice, and 0.5 ml of fresh medium was added to each well. Virus aliquots from each time point were titrated side-by-side on 24-well plates of Vero cells as described above.

### SDS-PAGE and Western blot analysis

Six-well plates of Vero or A549 cells were inoculated using an MOI of 1 PFU per cell with B/HPIV3, B/HPIV3/S-2P, and B/HPIV3/S-6P and incubated at 32°C. 48 h post-inoculation, cells were washed once with cold PBS and lysed using 300 µl LDS lysis buffer (Thermo Fisher Scientific) containing NuPAGE reducing reagent (Thermo Fisher Scientific). Cell lysates were further passed through a QIAshredder column (Qiagen), heated for 10 min at 95°C and separated on 4–12% Bis-Tris NuPAGE gels (Thermo Fisher Scientific) in the presence of antioxidant (Thermo Fisher Scientific). The resolved proteins were transferred to polyvinylidene difluoride membranes which were blocked with blocking buffer (LiCor) for 1h at room temperature. Then, blocked membranes were incubated with a goat hyperimmune serum to SARS-CoV-2 S and a rabbit polyclonal hyperimmune sera against purified HPIV3 in blocking buffer overnight at 4°C. A mouse monoclonal antibody to GAPDH (Sigma) was included as a loading control. Membranes were incubated with infrared dye-labeled secondary antibodies (donkey anti-rabbit IgG IRDye 680, donkey anti-goat IgG IRDye 800 or IRDye 680, and donkey anti-mouse IgG IRDye 800, LiCor). Intensities of individual protein bands from the acquired images were quantified after background correction using Image Studio software (LiCor).

The protein composition of virus particles was also analyzed. To do so, viruses were grown on Vero cells and the supernatant was purified by centrifugation through 30%/60% sucrose gradients. The sucrose-purified virus preparations were gently pelleted by centrifugation to remove sucrose as described previously [34]. The protein concentration of the purified virus preparations was determined using a BCA protein assay kit (Thermo Fisher Scientific) prior to the addition of lysis buffer. One µg of protein per lane was used for SDS-PAGE and Western blotting.

### Replication, immunogenicity, and protective efficacy against SARS-CoV-2 challenge in hamsters

All animal studies were approved by the NIAID Animal Care and Use Committee. In Experiment #1, groups (n=28) of 5- to 6-week-old male golden Syrian hamsters (Envigo Laboratories, Frederick, MD) were anesthetized and inoculated intranasally with 100 µl of Leibovitz’s L-15 medium (Thermo Fisher Scientific) containing 5 log_10_ PFU of B/HPIV3, B/HPIV3/S-2P, or B/HPIV3/S-6P viruses. On days 3, 5, and 7 post-inoculation, 5 hamsters per group were euthanized by CO_2_ inhalation, and nasal turbinates and lungs were collected to evaluate virus replication. NT and lung tissue samples for histology were obtained from one and two additional hamsters per group on days 3 and 5, respectively. For quantification of virus replication, tissues were homogenized in Leibovitz’s L-15 medium, and clarified homogenates were analyzed by dual-staining immunoplaque assay on Vero cells as described above. On day 26 post-immunization, sera were collected from the remaining 10 animals per group to evaluate the immunogenicity of the vaccine candidates. HPIV3 vector-specific neutralizing antibodies were detected by a 60% plaque reduction neutralization test (PRNT_60_) [35] on Vero cells in 24-well plates using HPIV3 expressing the eGFP protein. To determine the serum neutralizing antibody response to SARS-CoV-2, two-fold dilutions of heat-inactivated hamster sera were tested in a microneutralization assay for the presence of antibodies that neutralized the replication of 100 TCID_50_ of SARS-CoV-2 in Vero E6 cells as previously described [17]. Serum IgG antibodies to a secreted form of the S-2P protein [23] or its RBD [36] were measured by ELISA [17], and serum IgA antibodies to the secreted form of S-2P were measured by europium ion-enhanced DELFIA-TRF assay (Perkin Elmer) following the supplier’s protocol.

In Experiment #2, groups (n=45) of 5- to 6-week-old male golden Syrian hamsters were immunized intranasally with 5 log_10_ PFU of B/HPIV3, B/HPIV3/S-2P, or B/HPIV3/S-6P as described above. Sera were obtained on days 26 or 27. Hamsters in each immunization group were randomly distributed to 3 challenge virus subgroups of 15 animals per subgroup, and challenged intranasally with 4.5 log_10_ TCID_50_ of SARS-CoV-2 WA1/2020, the USA/CA_CDC_5574/2020 isolate (lineage B.1.1.7/Alpha variant) or USA/MD-HP01542/2021 (lineage B.1.351/Beta variant) in 100 µl. Five hamsters from each challenge virus subgroup were euthanized by CO_2_ inhalation on days 3 and 5 after challenge, and tissues were collected to evaluate challenge virus replication. The presence of challenge virus in clarified tissue homogenates was evaluated later by TCID_50_ assay on Vero E6 cells. The remaining 5 hamsters from each challenge virus subgroup were euthanized on day 21 after challenge, and serum was collected for antibody analysis.

### Immunohistopathology analysis

Lung tissue samples from hamsters were fixed in 10% neutral buffered formalin, processed through a Leica ASP6025 tissue processor (Leica Biosystems), and embedded in paraffin. Five μm tissue sections were stained with hematoxylin and eosin for routine histopathology. For IHC evaluation, sections were deparaffinized and rehydrated. After epitope retrieval, sections were labeled with goat hyperimmune serum to anti-SARS-CoV-2 S (N25-154) at 1:1000, and rabbit polyclonal anti-HPIV3 serum [37] at 1:500. Chromogenic staining was carried out on the Bond RX platform (Leica Biosystems) according to manufacturer-supplied protocols. Detection with DAB chromogen was completed using the Bond Polymer Refine Detection kit (Leica Biosystems). The VisUCyte anti-goat horseadish peroxidase (HRP) polymer (R&D Systems) replaced the standard Leica anti-rabbit HRP polymer from the kit to bind the SARS-CoV-2 S goat antibodies. Slides were finally cleared through gradient alcohol and xylene washes prior to mounting. Sections were examined by a board-certified veterinary pathologist using an Olympus BX51 light microscope. Images were acquired using an Olympus DP73 camera.

### Lentivirus-based pseudotype virus neutralization assay

Single-round luciferase-expressing pseudoviruses were generated by co-transfecting plasmids encoding SARS-CoV-2 S of isolate B.1.617.2/Delta or B.1.1.529/Omicron, luciferase reporter (pHR’ CMV Luc), lentivirus backbone (pCMV ΔR8.2), and human transmembrane protease serine 2 (TMPRSS2) at a ratio of 1:20:20:0.3 into HEK293T/17 cells (ATCC CRL-11268) using transfection reagent LiFect293. Three days post transfection, supernatants were collected and centrifugated at 478 x g for 10 min to remove cell debris and filtered through a 0.45 μm filter.

Pseudoviruses were then aliquoted and titrated. For the antibody neutralization assay, 6-point, 5-fold dilution series were prepared in DMEM medium supplemented with 10% FBS, 1 % Pen/Strep and 3 µg/ml puromycin. Fifty µl of the diluted sera were mixed with 50 µl of diluted pseudoviruses in 96-well plates and incubated for 30 min at 37°C. Then, ten thousand of ACE2-expressing 293T-cells (293T-hACE2.MF stable cell line) per well were added. Three days later, cells were lysed with Bright-Glo™ luciferase assay substrate (Promega), and luciferase activity (relative light units, RLU) was evaluated. The percent neutralization was normalized relative to uninfected cells as 100% neutralization and cells infected with only pseudoviruses as 0% neutralization. Fifty percent inhibitory concentration (IC_50_) titers were determined using a log (agonist) vs. normalized response (variable slope) nonlinear function in Prism v9 (GraphPad).

### ACE2 binding inhibition assay

A 10-plex ACE2 binding inhibition assay was performed as an alternative to a BSL3 SARS-CoV-2 neutralization assay [SARS-CoV-2 Panel 25 and 27 (K15586U and K15609U, respectively), MesoScale Diagnostics (MSD)], following the manufacturer’s instructions. The panels contain 96-well, 10-spot plates coated with soluble ectodomains of spike proteins from the vaccine-matched SARS-CoV-2 (Wuhan-Hu1, identical amino acid sequence as WA1/2020) and VoCs (B.1.1.7/Alpha, B.1.351/Beta, B.1.617.2/Delta, B.1.1.529/Omicron BA.1, BA.1+S[R346K], BA.1+S[L452R], BA.2, BA.2+S[L452M], BA.2+S[L452R], BA.2.12.1, BA.3, BA.4, BA.5 sublineages) and a variant under monitoring (B.1.640.2). In brief, plates were blocked with MSD blocker A for 30 min and washed with MSD wash buffer. Heat-inactivated hamster sera were diluted 1:20 in diluent and added to duplicate wells. After a one-hour incubation, sulfo-tag labelled ACE2 was added to each well. All incubations were performed on a plate shaker at room temperature. Following a one-hour incubation, plates were washed, MSD GOLD electrochemiluminescence read buffer B was added, and chemiluminescence of bound sulfo-tag-ACE2 was detected using a Meso 1300 Quickplex reader. The average electrochemiluminescence signals in duplicate wells for each serum were determined, as well as the maximum electrochemiluminescence signals of ACE2/S protein binding in no-serum control wells. The ACE2 neutralizing activity of each serum is represented as percent inhibition relative to no-serum controls.

### RT-qPCR analysis of gene expression in lung tissue

Total RNA was extracted from 0.1 ml of lung homogenates (0.1 g/ml) using the TRIzol LS Reagent and Phasemaker Tubes Complete System (Thermo Fisher) along with the PureLink RNA Mini Kit (Thermo Fisher) following the manufacturer’s instructions, and eluted in 0.1 ml RNase-free water. Total RNA was also extracted from lung homogenates of five control hamsters (non-immunized and non-challenged) in the same manner. cDNA was synthesized from 7 µl of RNA using the High-Capacity RNA-to-cDNA Kit (Thermo Fisher). TaqMan assays (Thermo Fisher) specific for hamster (*Mesocricetus auratus*) CXCL10, Mx2, or IFN-L [38, 39] were performed using the TaqMan Fast Advanced Master Mix (Thermo Fisher), and hamster beta-actin was included as a housekeeping gene. qPCR results were analyzed using the comparative threshold cycle (ΔΔC_T_) method, normalized to beta-actin, and expressed as fold changes over the average expression of five uninfected, unchallenged hamsters.

To detect viral genomic N (gN), E (gE) RNA, and subgenomic E (sgE) mRNA of SARS-CoV-2 challenge virus WA1/2020, B.1.1.7/Alpha, or B.1.351/Beta in the lung homogenates, cDNA was synthesized from 8 μl of total RNA extracted as described above, and TaqMan qPCRs for genomic N, E, and sgE were performed using previously-described primers and probes [40–42] and the TaqMan RNA-to-Ct 1-Step Kit (Thermo Fisher). Assays were performed on a QuantStudio 7 Pro real-time PCR system (Thermo Fisher). Standard curves were generated using serially diluted pcDNA3.1 plasmids containing gN, gE, or sgE sequences. The sensitivity of the TaqMan assay was 200 copies, corresponding to a limit of detection of 5.4 log_10_ copies per g of tissue.

### Statistical analysis

Data sets were assessed for significance using one-way ANOVA with Tukey’s or Sidak multiple comparisons test, two-way ANOVA with Tukey post-hoc test or Mixed-effects analysis with Tukey post-hoc test using Prism 9 (GraphPad Software). Data were only considered significant at P ≤ 0.05. Details on the statistical comparisons can be found in the figures, figure legends, and results. Values of n are provided in the text and in the figure legends.

## ACKNOWLEDGEMENTS

The authors thank the staff of the NIAID Comparative Medicine Branch for animal study support. We thank Barney Graham and Kizzmekia Corbett, Vaccine Research Center, NIAID, NIH, and Jason McLellan, University of Texas, Austin, TX, for providing plasmid 2019-nCoV S-2P_dFurin_F3CH2S encoding the prefusion-stabilized version of the SARS-CoV-2 S ectodomain which was used to generate ELISA antigen for our studies. The SARS-CoV-2 S protein containing the receptor-binding domain (RBD) was obtained from David Veesler through BEI Resources, NIAID, NIH. We thank Ruturaj Masvekar and Bibi Bielekova, Laboratory of Clinical Immunology & Microbiology, NIAID, NIH for their help with the MSD plate reader. This research was supported by the Intramural Research Program of the NIAID, NIH (Project number ZIA AI001298-01).

## AUTHOR CONTRIBUTIONS

Conceptualization: H.-S.P., X.L., Y.M., C.LN, and U.J.B.. Methodology, formal analysis, visualization: H.-S.P., X.L., Y.M., C.S., L.Y., C.L., I.N.M., P.Z., P.L., C.LN and U.J.B.. Investigation: H.-S.P., X.L., Y.M., C.S., L.Y., C.L., I.N.M., P.Z., P.L., and C.LN.. Resources: R.J., N.L.G., and S.M.B.. Writing – Original Draft: X.L., H.-S.P., C.LN. and U.J.B.. Writing – Review and Editing: All authors.

## FIGURE CAPTIONS

## SUPPLEMENTAL FIGURE CAPTIONS

**Figure S1.**
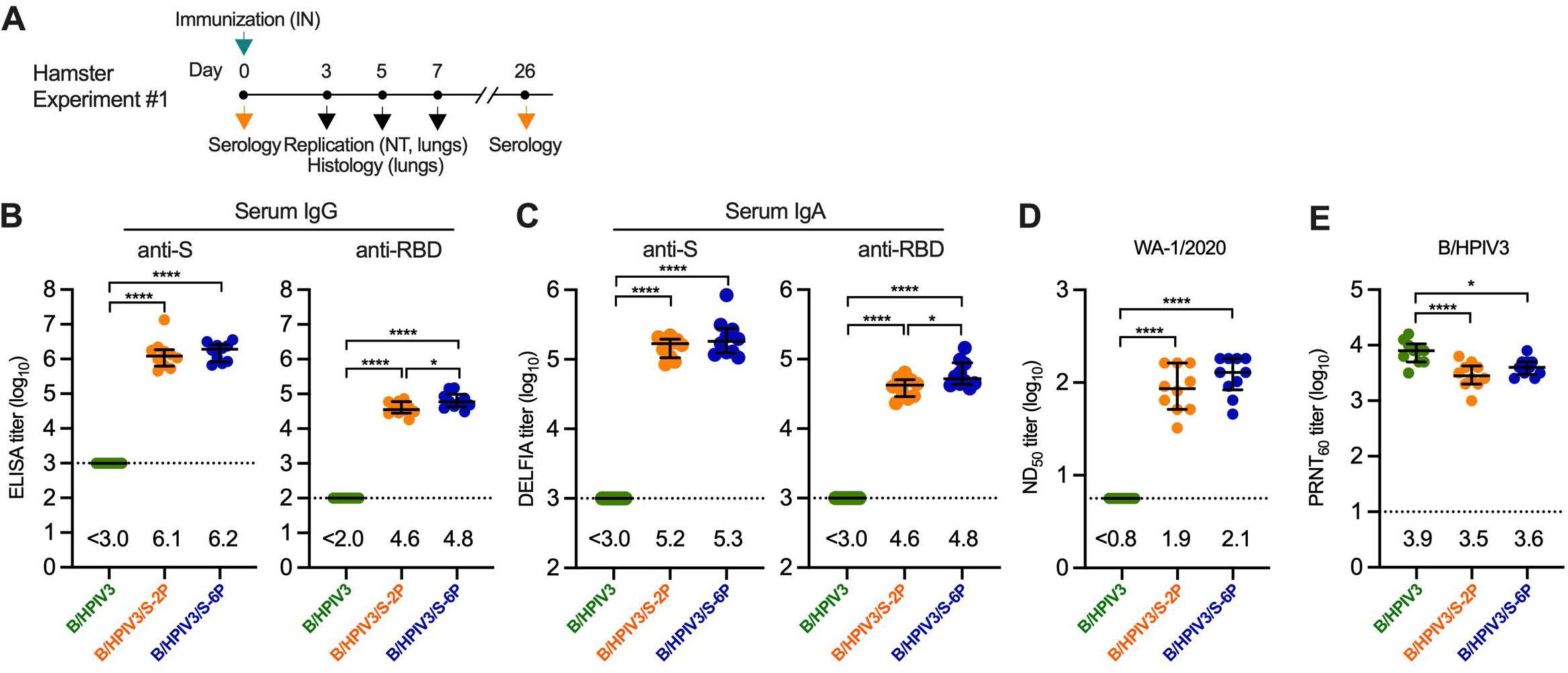
Immunogenicity of B/HPIV3, B/HPIV3/S-2P, or B/HPIV3/S-6P in hamsters (Additional data from Experiment 1, related to Figure 2) (A) Schematic overview of Experiment 1. Hamsters were immunized intranasally with 5.0 log_10_ PFU of B/HPIV3, B/HPIV3/S-2P, or B/HPIV3/S-6P. Vaccine virus titers detected in respiratory tissues on days 3, 5, and 7 after immunization are shown in Figure 2. Sera from n=10 animals per group were collected on day 26 after immunization. (B) IgG ELISA titers to a secreted form of the S-2P protein or to a fragment of the S protein (aa 328-531) containing SARS-CoV-2 receptor-binding domain (RBD) and (C) IgA titers to S-2P or the RBD, determined by dissociation-enhanced lanthanide time-resolved fluorescence (DELFIA-TRF) assay. (D) The 50% SARS-CoV-2 neutralizing titers (ND_50_) were determined on Vero E6 cells against the vaccine-matched strain WA1/2020. (E) Sera were also analyzed to determine the 60% plaque reduction neutralization titers (PRNT_60_) to HPIV3. Each hamster is represented by a symbol and GMTs and standard deviations indicated. For (B) and (E), GMTs are indicated above the x axis. The limit of detection is indicated by a dashed line. Medians and interquartile ranges are shown *=P<0.05; ****=P<0.0001 (One-way ANOVA with Tukey multiple comparisons).

**Figure S2.**
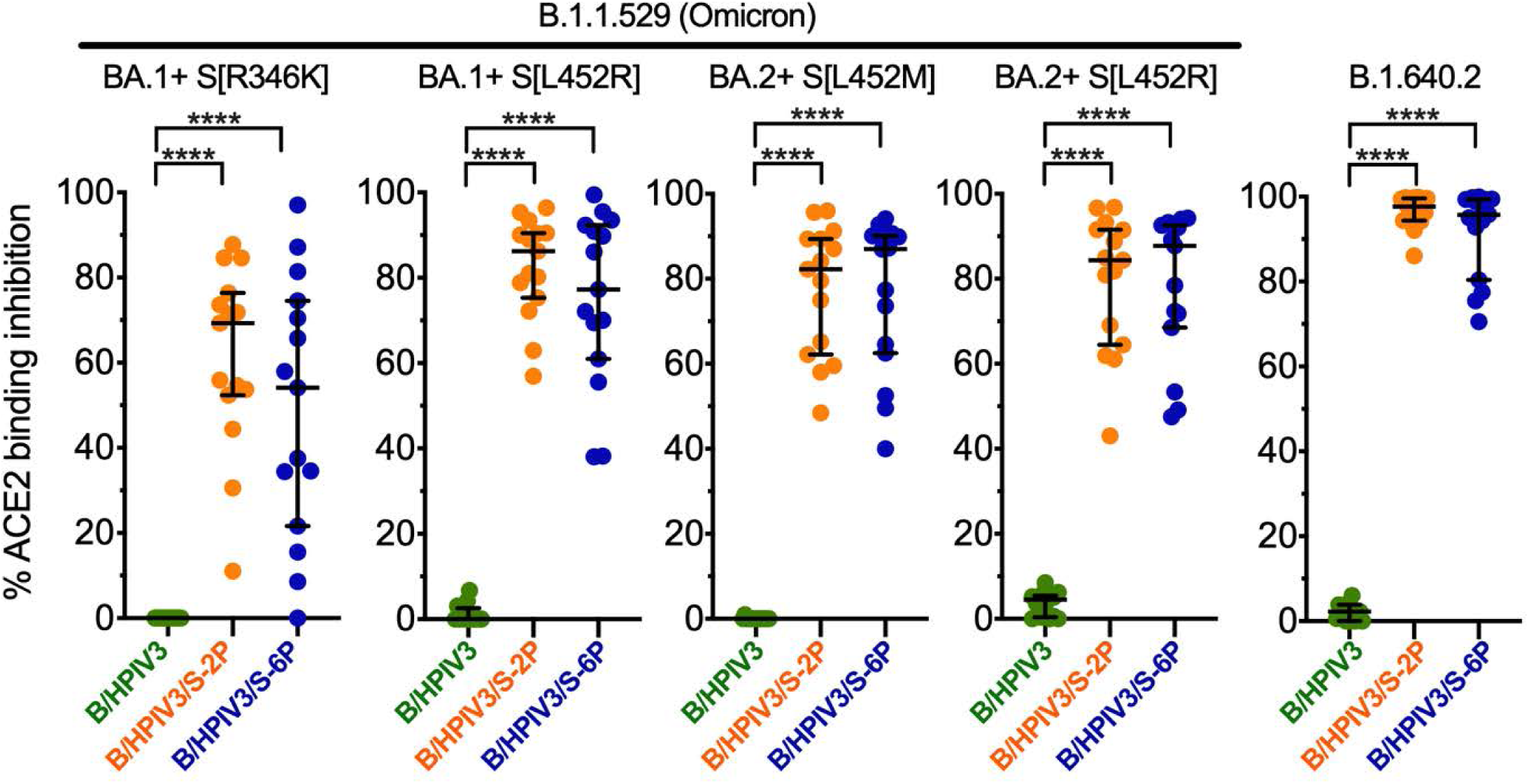
Inhibition by serum antibodies from B/HPIV3/S-2P- and B/HPIV3/S-6P-immunized hamsters of ACE2 binding to S proteins of B.1.1.529 and B.1.640 variants, related to Figure 3G. An ACE2 binding inhibition assay was used as an alternative for a BSL3 live-virus neutralization assay. Heat-inactivated hamster sera were diluted 1:20 and added to duplicate wells of 96-well plates spot-coated with the indicated S proteins. The percent binding inhibition of sulfo-tag labelled ACE2 to S proteins of the indicated VoCs by serum antibodies from immunized hamsters was determined by electrochemiluminescence. Each hamster is represented by a symbol, and medians and interquartile ranges are shown. ****=P<0.0001 (One-way ANOVA with Tukey multiple comparisons).

**Figure S3.**
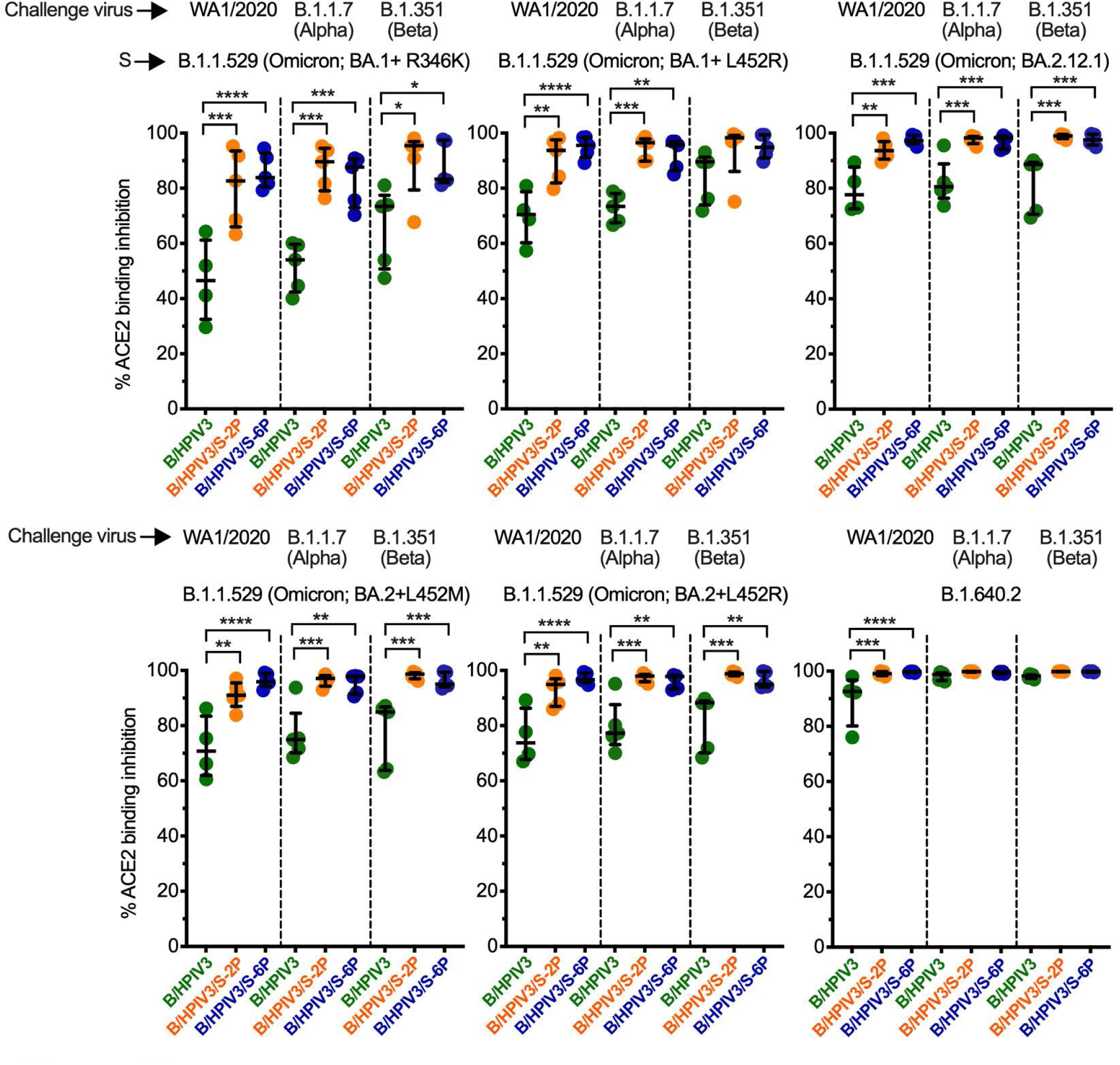
Binding inhibition of S to ACE2 by the sera of B/HPIV3/S-2P- and B/HPIV3/S-6P-immunized hamsters three weeks after challenge with SARS-CoV-2 WA1/2020 or VoCs B.1.1.7/Alpha or B.1.351/Beta, related to Figure 6B. An ACE2 binding inhibition assay was used as an alternative for a BSL3 live-virus neutralization assay. Heat-inactivated hamster sera were diluted 1:20 and added to duplicate wells of 96-well plates spot-coated with the indicated S proteins. The percent binding inhibition of sulfo-tag labelled ACE2 to S proteins of the indicated VoCs by serum antibodies from immunized hamsters was determined by electrochemiluminescence. Each hamster is represented by a symbol, and medians and interquartile ranges are shown. *=P<0.05; **=P<0.01; ***=P<0.001; ****=P<0.0001 (One-way ANOVA with Sidák’s multiple comparisons).

## REFERENCES

1. Martin B, DeWitt PE, Russell S, Anand A, Bradwell KR, Bremer C, et al. Characteristics, Outcomes, and Severity Risk Factors Associated With SARS-CoV-2 Infection Among Children in the US National COVID Cohort Collaborative. JAMA Netw Open. 2022;5(2):e2143151. Epub 20220201. doi: 10.1001/jamanetworkopen.2021.43151. PubMed PMID: 35133437; PubMed Central PMCID: PMCPMC8826172.

2. Cui X, Zhao Z, Zhang T, Guo W, Guo W, Zheng J, et al. A systematic review and meta-analysis of children with coronavirus disease 2019 (COVID-19). J Med Virol. 2021;93(2):1057–69. Epub 20200928. doi: 10.1002/jmv.26398. PubMed PMID: 32761898; PubMed Central PMCID: PMCPMC7436402.

3. Zachariah P, Johnson CL, Halabi KC, Ahn D, Sen AI, Fischer A, et al. Epidemiology, Clinical Features, and Disease Severity in Patients With Coronavirus Disease 2019 (COVID-19) in a Children’s Hospital in New York City, New York. JAMA Pediatr. 2020;174(10):e202430. Epub 20201005. doi: 10.1001/jamapediatrics.2020.2430. PubMed PMID: 32492092; PubMed Central PMCID: PMCPMC7270880.

4. Anderson EJ, Rouphael NG, Widge AT, Jackson LA, Roberts PC, Makhene M, et al. Safety and Immunogenicity of SARS-CoV-2 mRNA-1273 Vaccine in Older Adults. N Engl J Med. 2020;383(25):2427–38. Epub 20200929. doi: 10.1056/NEJMoa2028436. PubMed PMID: 32991794; PubMed Central PMCID: PMCPMC7556339.

5. Ewer KJ, Barrett JR, Belij-Rammerstorfer S, Sharpe H, Makinson R, Morter R, et al. T cell and antibody responses induced by a single dose of ChAdOx1 nCoV-19 (AZD1222) vaccine in a phase 1/2 clinical trial. Nat Med. 2021;27(2):270–8. Epub 20201217. doi: 10.1038/s41591-020-01194-5. PubMed PMID: 33335323.

6. Sahin U, Muik A, Vogler I, Derhovanessian E, Kranz LM, Vormehr M, et al. BNT162b2 vaccine induces neutralizing antibodies and poly-specific T cells in humans. Nature. 2021;595(7868):572–7. Epub 20210527. doi: 10.1038/s41586-021-03653-6. PubMed PMID: 34044428.

7. Lund FE, Randall TD. Scent of a vaccine. Science. 2021;373(6553):397–9. doi: 10.1126/science.abg9857. PubMed PMID: 34437109.

8. Karron RA, Thumar B, Schappell E, Surman S, Murphy BR, Collins PL, et al. Evaluation of two chimeric bovine-human parainfluenza virus type 3 vaccines in infants and young children. Vaccine. 2012;30(26):3975–81. Epub 2011/12/20. doi: 10.1016/j.vaccine.2011.12.022. PubMed PMID: 22178099; PubMed Central PMCID: PMCPMC3509782.

9. Schmidt AC, McAuliffe JM, Huang A, Surman SR, Bailly JE, Elkins WR, et al. Bovine parainfluenza virus type 3 (BPIV3) fusion and hemagglutinin-neuraminidase glycoproteins make an important contribution to the restricted replication of BPIV3 in primates. J Virol. 2000;74(19):8922–9. Epub 2000/09/12. doi: 10.1128/jvi.74.19.8922-8929.2000. PubMed PMID: 10982335; PubMed Central PMCID: PMCPMC102087.

10. Howard LM, Edwards KM, Zhu Y, Williams DJ, Self WH, Jain S, et al. Parainfluenza Virus Types 1-3 Infections Among Children and Adults Hospitalized With Community-acquired Pneumonia. Clin Infect Dis. 2021;73(11):e4433–e43. doi: 10.1093/cid/ciaa973. PubMed PMID: 32681645; PubMed Central PMCID: PMCPMC8662767.

11. DeGroote NP, Haynes AK, Taylor C, Killerby ME, Dahl RM, Mustaquim D, et al. Human parainfluenza virus circulation, United States, 2011-2019. J Clin Virol. 2020;124:104261. Epub 20200109. doi: 10.1016/j.jcv.2020.104261. PubMed PMID: 31954277; PubMed Central PMCID: PMCPMC7106518.

12. Li Y, Reeves RM, Wang X, Bassat Q, Brooks WA, Cohen C, et al. Global patterns in monthly activity of influenza virus, respiratory syncytial virus, parainfluenza virus, and metapneumovirus: a systematic analysis. Lancet Glob Health. 2019;7(8):e1031–e45. doi: 10.1016/S2214-109X(19)30264-5. PubMed PMID: 31303294.

13. van Wyke Coelingh KL, Winter CC, Tierney EL, London WT, Murphy BR. Attenuation of bovine parainfluenza virus type 3 in nonhuman primates and its ability to confer immunity to human parainfluenza virus type 3 challenge. J Infect Dis. 1988;157(4):655–62. doi: 10.1093/infdis/157.4.655. PubMed PMID: 2831282.

14. Schmidt AC, McAuliffe JM, Murphy BR, Collins PL. Recombinant bovine/human parainfluenza virus type 3 (B/HPIV3) expressing the respiratory syncytial virus (RSV) G and F proteins can be used to achieve simultaneous mucosal immunization against RSV and HPIV3. J Virol. 2001;75(10):4594–603. doi: 10.1128/JVI.75.10.4594-4603.2001. PubMed PMID: 11312329; PubMed Central PMCID: PMCPMC114212.

15. Bernstein DI, Malkin E, Abughali N, Falloon J, Yi T, Dubovsky F, et al. Phase 1 study of the safety and immunogenicity of a live, attenuated respiratory syncytial virus and parainfluenza virus type 3 vaccine in seronegative children. Pediatr Infect Dis J. 2012;31(2):109–14. Epub 2011/09/20. doi: 10.1097/INF.0b013e31823386f1. PubMed PMID: 21926667.

16. Liang B, Ngwuta JO, Herbert R, Swerczek J, Dorward DW, Amaro-Carambot E, et al. Packaging and Prefusion Stabilization Separately and Additively Increase the Quantity and Quality of Respiratory Syncytial Virus (RSV)-Neutralizing Antibodies Induced by an RSV Fusion Protein Expressed by a Parainfluenza Virus Vector. J Virol. 2016;90(21):10022–38. Epub 20161014. doi: 10.1128/JVI.01196-16. PubMed PMID: 27581977; PubMed Central PMCID: PMCPMC5068507.

17. Liu X, Luongo C, Matsuoka Y, Park HS, Santos C, Yang L, et al. A single intranasal dose of a live-attenuated parainfluenza virus-vectored SARS-CoV-2 vaccine is protective in hamsters. Proc Natl Acad Sci U S A. 2021;118(50). Epub 2021/12/09. doi: 10.1073/pnas.2109744118. PubMed PMID: 34876520.

18. Chan JF, Zhang AJ, Yuan S, Poon VK, Chan CC, Lee AC, et al. Simulation of the Clinical and Pathological Manifestations of Coronavirus Disease 2019 (COVID-19) in a Golden Syrian Hamster Model: Implications for Disease Pathogenesis and Transmissibility. Clin Infect Dis. 2020;71(9):2428–46. doi: 10.1093/cid/ciaa325. PubMed PMID: 32215622; PubMed Central PMCID: PMCPMC7184405.

19. Imai M, Iwatsuki-Horimoto K, Hatta M, Loeber S, Halfmann PJ, Nakajima N, et al. Syrian hamsters as a small animal model for SARS-CoV-2 infection and countermeasure development. Proc Natl Acad Sci U S A. 2020;117(28):16587–95. Epub 20200622. doi: 10.1073/pnas.2009799117. PubMed PMID: 32571934; PubMed Central PMCID: PMCPMC7368255.

20. Sia SF, Yan LM, Chin AWH, Fung K, Choy KT, Wong AYL, et al. Pathogenesis and transmission of SARS-CoV-2 in golden hamsters. Nature. 2020;583(7818):834–8. Epub 20200514. doi: 10.1038/s41586-020-2342-5. PubMed PMID: 32408338; PubMed Central PMCID: PMCPMC7394720.

21. Hsieh CL, Goldsmith JA, Schaub JM, DiVenere AM, Kuo HC, Javanmardi K, et al. Structure-based design of prefusion-stabilized SARS-CoV-2 spikes. Science. 2020;369(6510):1501–5. Epub 2020/07/25. doi: 10.1126/science.abd0826. PubMed PMID: 32703906; PubMed Central PMCID: PMCPMC7402631.

22. Le Nouen C, Nelson CE, Liu X, Park HS, Matsuoka Y, Luongo C, et al. Intranasal pediatric parainfluenza virus-vectored SARS-CoV-2 vaccine is protective in monkeys. Cell. 2022. Epub 20221110. doi: 10.1016/j.cell.2022.11.006. PubMed PMID: 36423629.

23. Wrapp D, Wang N, Corbett KS, Goldsmith JA, Hsieh CL, Abiona O, et al. Cryo-EM structure of the 2019-nCoV spike in the prefusion conformation. Science. 2020;367(6483):1260–3. Epub 2020/02/23. doi: 10.1126/science.abb2507. PubMed PMID: 32075877; PubMed Central PMCID: PMCPMC7164637.

24. Yang Y, Shen C, Li J, Yuan J, Wei J, Huang F, et al. Plasma IP-10 and MCP-3 levels are highly associated with disease severity and predict the progression of COVID-19. J Allergy Clin Immunol. 2020;146(1):119–27 e4. Epub 20200429. doi: 10.1016/j.jaci.2020.04.027. PubMed PMID: 32360286; PubMed Central PMCID: PMCPMC7189843.

25. Pegu A, O’Connell SE, Schmidt SD, O’Dell S, Talana CA, Lai L, et al. Durability of mRNA-1273 vaccine-induced antibodies against SARS-CoV-2 variants. Science. 2021;373(6561):1372–7. Epub 20210813. doi: 10.1126/science.abj4176. PubMed PMID: 34385356; PubMed Central PMCID: PMCPMC8691522.

26. Doria-Rose N, Suthar MS, Makowski M, O’Connell S, McDermott AB, Flach B, et al. Antibody Persistence through 6 Months after the Second Dose of mRNA-1273 Vaccine for Covid-19. N Engl J Med. 2021;384(23):2259–61. Epub 20210406. doi: 10.1056/NEJMc2103916. PubMed PMID: 33822494; PubMed Central PMCID: PMCPMC8524784.

27. Doria-Rose NA, Shen X, Schmidt SD, O’Dell S, McDanal C, Feng W, et al. Booster of mRNA-1273 Strengthens SARS-CoV-2 Omicron Neutralization. medRxiv. 2021. Epub 20211220. doi: 10.1101/2021.12.15.21267805. PubMed PMID: 34931200; PubMed Central PMCID: PMCPMC8687471.

28. Chemaitelly H, Ayoub HH, AlMukdad S, Coyle P, Tang P, Yassine HM, et al. Duration of mRNA vaccine protection against SARS-CoV-2 Omicron BA.1 and BA.2 subvariants in Qatar. Nature communications. 2022;13(1):3082. Epub 20220602. doi: 10.1038/s41467-022-30895-3. PubMed PMID: 35654888; PubMed Central PMCID: PMCPMC9163167.

29. Liang B, Ngwuta JO, Surman S, Kabatova B, Liu X, Lingemann M, et al. Improved Prefusion Stability, Optimized Codon Usage, and Augmented Virion Packaging Enhance the Immunogenicity of Respiratory Syncytial Virus Fusion Protein in a Vectored-Vaccine Candidate. J Virol. 2017;91(15). Epub 20170712. doi: 10.1128/JVI.00189-17. PubMed PMID: 28539444; PubMed Central PMCID: PMCPMC5651718.

30. Tostanoski LH, Yu J, Mercado NB, McMahan K, Jacob-Dolan C, Martinot AJ, et al. Immunity elicited by natural infection or Ad26.COV2.S vaccination protects hamsters against SARS-CoV-2 variants of concern. Sci Transl Med. 2021;13(618):eabj3789. Epub 20211103. doi: 10.1126/scitranslmed.abj3789. PubMed PMID: 34705477; PubMed Central PMCID: PMCPMC8818312.

31. Hawman DW, Meade-White K, Archer J, Leventhal SS, Wilson D, Shaia C, et al. SARS-CoV2 variant-specific replicating RNA vaccines protect from disease following challenge with heterologous variants of concern. Elife. 2022;11. Epub 20220222. doi: 10.7554/eLife.75537. PubMed PMID: 35191378; PubMed Central PMCID: PMCPMC8983041.

32. Fischer RJ, van Doremalen N, Adney DR, Yinda CK, Port JR, Holbrook MG, et al. ChAdOx1 nCoV-19 (AZD1222) protects Syrian hamsters against SARS-CoV-2 B.1.351 and B.1.1.7. Nature communications. 2021;12(1):5868. Epub 20211007. doi: 10.1038/s41467-021-26178-y. PubMed PMID: 34620866; PubMed Central PMCID: PMCPMC8497486.

33. Subbarao K, McAuliffe J, Vogel L, Fahle G, Fischer S, Tatti K, et al. Prior infection and passive transfer of neutralizing antibody prevent replication of severe acute respiratory syndrome coronavirus in the respiratory tract of mice. J Virol. 2004;78(7):3572–7. Epub 2004/03/16. PubMed PMID: 15016880; PubMed Central PMCID: PMCPMC371090.

34. Munir S, Le Nouen C, Luongo C, Buchholz UJ, Collins PL, Bukreyev A. Nonstructural proteins 1 and 2 of respiratory syncytial virus suppress maturation of human dendritic cells. J Virol. 2008;82(17):8780–96. Epub 2008/06/20. doi: 10.1128/JVI.00630-08. PubMed PMID: 18562519.

35. Liang B, Surman S, Amaro-Carambot E, Kabatova B, Mackow N, Lingemann M, et al. Enhanced Neutralizing Antibody Response Induced by Respiratory Syncytial Virus Pre-fusion F Protein Expressed by a Vaccine Candidate. J Virol. 2015. doi: 10.1128/JVI.01373-15. PubMed PMID: 26157122.

36. Walls AC, Park YJ, Tortorici MA, Wall A, McGuire AT, Veesler D. Structure, Function, and Antigenicity of the SARS-CoV-2 Spike Glycoprotein. Cell. 2020;183(6):1735. doi: 10.1016/j.cell.2020.11.032. PubMed PMID: 33306958; PubMed Central PMCID: PMCPMC7833104.

37. Liang B, Munir S, Amaro-Carambot E, Surman S, Mackow N, Yang L, et al. Chimeric bovine/human parainfluenza virus type 3 expressing respiratory syncytial virus (RSV) F glycoprotein: effect of insert position on expression, replication, immunogenicity, stability, and protection against RSV infection. J Virol. 2014;88(8):4237–50. doi: 10.1128/JVI.03481-13. PubMed PMID: 24478424; PubMed Central PMCID: PMCPMC3993740.

38. Zivcec M, Safronetz D, Haddock E, Feldmann H, Ebihara H. Validation of assays to monitor immune responses in the Syrian golden hamster (Mesocricetus auratus). J Immunol Methods. 2011;368(1-2):24–35. Epub 20110217. doi: 10.1016/j.jim.2011.02.004. PubMed PMID: 21334343; PubMed Central PMCID: PMCPMC3085612.

39. Sanchez-Felipe L, Vercruysse T, Sharma S, Ma J, Lemmens V, Van Looveren D, et al. A single-dose live-attenuated YF17D-vectored SARS-CoV-2 vaccine candidate. Nature. 2021;590(7845):320–5. Epub 20201201. doi: 10.1038/s41586-020-3035-9. PubMed PMID: 33260195.

40. Wolfel R, Corman VM, Guggemos W, Seilmaier M, Zange S, Muller MA, et al. Virological assessment of hospitalized patients with COVID-2019. Nature. 2020;581(7809):465–9. Epub 2020/04/03. doi: 10.1038/s41586-020-2196-x. PubMed PMID: 32235945.

41. Corman VM, Landt O, Kaiser M, Molenkamp R, Meijer A, Chu DK, et al. Detection of 2019 novel coronavirus (2019-nCoV) by real-time RT-PCR. Euro surveillance : bulletin Europeen sur les maladies transmissibles = European communicable disease bulletin. 2020;25(3). Epub 2020/01/30. doi: 10.2807/1560-7917.ES.2020.25.3.2000045. PubMed PMID: 31992387; PubMed Central PMCID: PMCPMC6988269.

42. Chandrashekar A, Liu J, Martinot AJ, McMahan K, Mercado NB, Peter L, et al. SARS-CoV-2 infection protects against rechallenge in rhesus macaques. Science. 2020;369(6505):812-Epub 2020/05/22. doi: 10.1126/science.abc4776. PubMed PMID: 32434946; PubMed Central PMCID: PMCPMC7243369.

